# Integration of multi-level dental diversity links macro-evolutionary patterns to ecological strategies across sharks

**DOI:** 10.1101/2024.10.30.621069

**Authors:** Roland Zimm, Vitória Tobias-Santos, Nicolas Goudemand

## Abstract

Sharks are notorious for their exceptional dental diversity, which is frequently used as a proxy for ecological function. However, functional inferences from morphology need to consider morphological features across different organizational scales, from small cusplets to patterns at the level of the entire jaw. Here, we deploy a set of classic and novel morphometric approaches to quantify morphological features ranging from sub-dental features to whole dentitions within a large ensemble of species encompassing all extant orders of sharks. We then correlate these measures with habitat, feeding and body size traits and track their variation as a function of genetic distance, a measure of trait adaptability. Intriguingly, sharks tend to either explore tooth-level or dentition-level complexity, resulting in two distinct groups with key differences in tooth symmetry, graduality of heterodont change, and depth of habitat. Overall, we find that intermediate levels of resolution, namely monognathic heterodonty in comparison with dignathic heterodonty and tooth-level shape descriptors, show the strongest predictive power for ecological traits, while exhibiting low phylogenetic signal, which suggests a more dynamic adaptability on shorter evolutionary timescales. This raises macro-evolutionary interpretations about the evolvability of nested modular phenotypic structures, with likely important implications for paleo-ecological inferences from sequentially homologous traits.

## Introduction

Teeth have been used as a powerful proxy for ecological function and adaptive evolution across vertebrates^1–7^As a hyper-diverse structure displaying considerable morphological change within short evolutionary frameworks^8–10^, tooth shape tends to be fine-tuned for food acquisition and mastication strategies^6^. This is particularly important for the reconstruction and study of past ecosystems where fossil teeth are often among the most abundant remnants within an inherently patchy and limited data record, providing critical information about ecological niche occupancy^2,4,11–13^While complexity and form of isolated teeth are informative about potential functions^14–17^, teeth tend to act together as a whole (or partial) dentition, forming an emergent functional unit. Thus, single-tooth morphology is only one of several different organizational levels - spanning from sub-dental features (e.g.serrations) to entire dentitions - that matter for specific functional aspects and their integration. This amounts to a limitation in paleontological studies often relying on dispersed isolated teeth whose relative positions are deduced indirectly^18^.

The functional integration of teeth, as whole dentitions or by regional subfunctionalization, is further illustrated by the widespread occurrence of heterodonty (juxtaposition of differently-shaped teeth). Like mammals, sharks exhibit conspicuous tooth morphological variation at different scales, from single tooth to dentition levels. Interestingly, this is not a recently evolved pattern, as heterodont arrangements of multicuspid teeth are described among the earliest sharks^19^.Albeit less studied as mammals, shark odontogenesis shares many central features with the former class, involving conserved regulatory pathways and developmental mechanisms^20–22^, besides notable differences^20,23^. Such deep similarities across long phylogenetic distances point towards a kernel of developmental mechanisms capable of generating highly diverse dental morphologies both between and within individuals, which has been explored experimentally and computationally^23–25^. While a system of tooth classes is well-established in mammals^26,27^, systematic knowledge about heterodonty biases in sharks is sparse. For specific low-rank taxonomic groups, patterns of tooth shape and size variation along the jaw are often diagnostic^28,29^. Some of these patterns have been linked to feeding mechanics, emphasizing the importance of dentition-level perspectives when connecting morphology and ecological functions^30^, while stark tooth morphology differences persist between sexes and age cohorts^29,31–34^. This is why several studies have dissected dentitions morphometrically within proximate phylogenetic contexts^31,33,35^.

Complementarily, many studies describing shark tooth morphospaces have used isolated teeth across wider taxonomic levels, identifying clade- and ecotype-specific clusters and distributions^33,36–39^.Representing higher organizational levels, jaw geometry, cranial shape, musculature and other anatomic macro-features have been connected to different feeding strategies^40–42^, underlining the adaptive interplay of traits on different scales. This suggests that a class-wide analysis of functional associations between dental variation and complexity at different scales and ecological functions might be highly informative. Here, we elucidate such macro-patterns in the light of environmental and life-history traits, applying a novel combination of morphometric tools. This may critically contribute to understanding which organizational level, across an entire vertebrate class, is most relevant or predictive for functional traits, a significant question across ecology, paleontology and evolution.

## Results

### 1. Heterodonty is widespread across sharks

Heterodonty reflects the degree of tooth shape variation within a given individual. Such variation is not always subtle and gradual, limiting the use of common homology-based morphometrics tools (e.g.landmark and semi-landmark-based approaches), arguably biasing efforts to focus on species with gradual heterodonty^29,31^and making class-level comparisons rare^36,37^. We calculate different types of within-toothrow heterodonty, namely differences between neighbouring teeth within the same jaw (sequential monognathic heterodonty, HMS), between all teeth within the same jaw (total monognatic heterodonty, HMT), and between teeth from the same relative positions in opposing jaws (dignathic heterodonty, HDG) (cf.FIG.1). This pipeline allows reducing complex and multidimensional features to one-dimensional measures, and comparing morphologically heterogeneous taxa. Since heterodonty measures quantify shape variation between units, they can be considered a proxy for high-level (i.e.jaw-level) complexity. We also devise different proxies for single-tooth morphological complexity based on 2D-outline characteristics (see FIG.11), enabling us to contrast tooth and dentition-level features.

**Figure 1.**
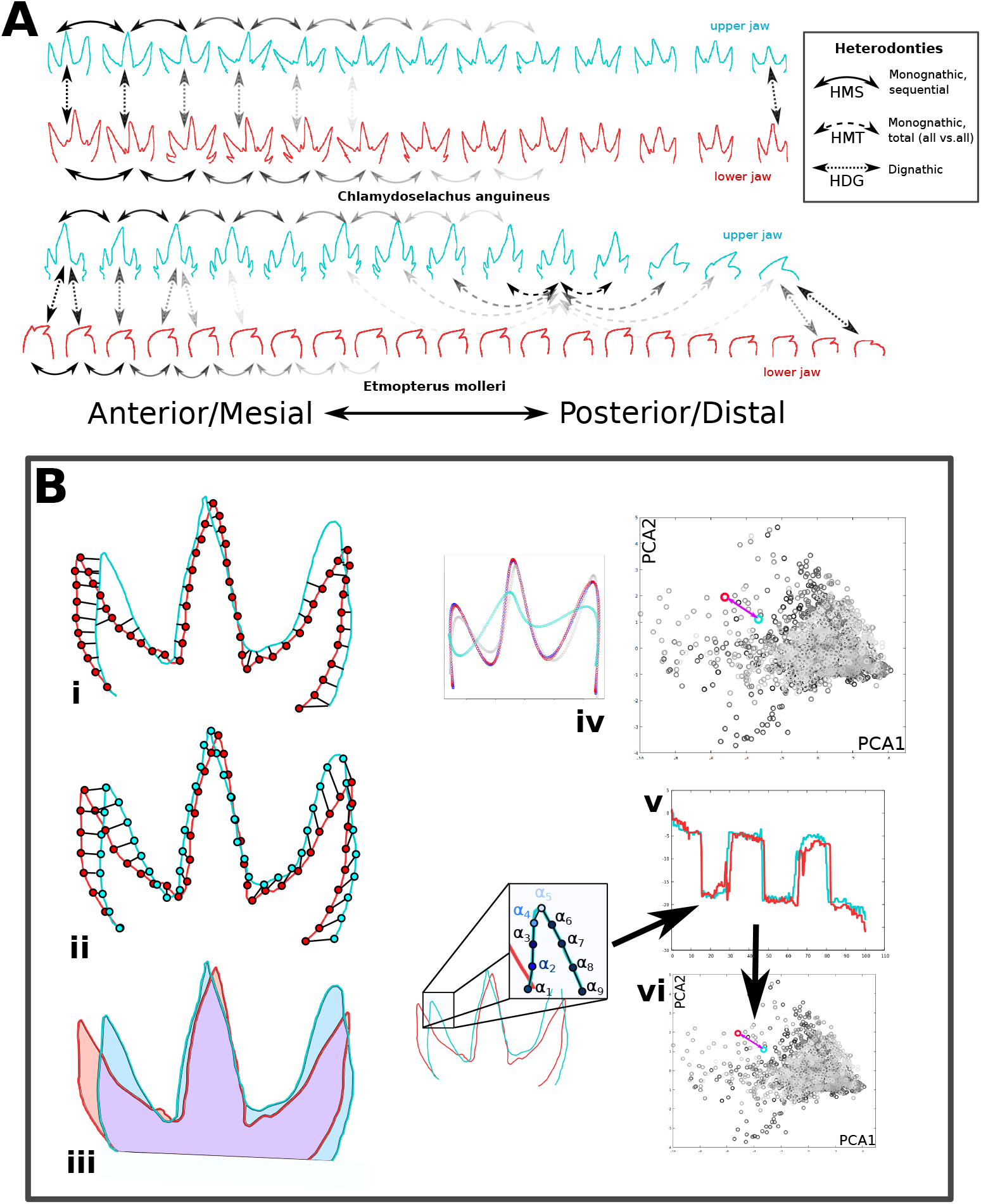
Overview of heterodonty measures. **A)** Different types of heterodonty: Heterodonty, a dentition-level disparity measure, can refer to (1) average differences between successive teeth: HMS, (2) differences between pairs of teeth belonging to opposing jaws, also termed dignathic heterodonty: HDG, (3) differences between all teeth within the same jaw: HMT. Here, these different measures are exemplified using differently dashed arrows. While the upper dentition of *C*.*anguineus* shows fairly similar teeth between upper (blue) and lower(red) jaw, the lower dentition of *E*.*molleri* displays conspicuous dignathic heterodonty. **B)** Comparing tooth similarity: Lacking a universally accepted metric of phenotypic difference, we deployed 6 different measures to quantify differences between two given tooth outlines: (i) Euclidean mean distance; the average distance between equally distant points along the tooth outline and the closest points of the second outline, respectively. Note that the measure is only shown for distances from given points on the red onto the blue outline, albeit being actually applied both ways. (ii) Homologous outline distance: we average pairwise distances between a sequence of equally spaced points on the two outlines, i.e. we compare points with the same numbers/indices. (iii) Superimposed area overlap: similarity is calculated as the ratio between overlapping to non-overlapping area. (iv) Discrete Cosine Fourier distance: The discrete cosine Fourier describing outline shapes yields a number of coefficients defining a sequence of cosine functions with increasing harmonics. Euclidean distances between corresponding coefficients are used to quantify shape distance. (v) Outline angle distance: Assuming that surface angles reflect relevant outline features, we calculate the distance between the functions summing outline angles at an intermediate resolution of outline points (i.e.for 100 outline points each) for two outlines. (vi) Angle Function Discrete Cosine Fourier: We apply the discrete cosine Fourier as in (iv) on the functions used in (v), in an analogous manner. Note that superpositions of the tooth outlines required for (i-iii) are anteceded by a partial Procrustes alignment.

In order to assess the prevalence of heterodonty across sharks, we collected 2D shape information about complete or nearly complete dentitions from 51 species, using an open data collection (J-elasmo^43^), representing all extant shark orders. The above defined measures identified substantial levels of heterodonty within all major clades of sharks, albeit in varying degrees (FIG.2). While no significant difference emerges between the two superorders Squalomorphii and Galeomorphii regarding total and maximal monognathic heterodonty (P_HMT_=0.3386, P_HMTmax_=0.928, two-sided Wilcoxon test), sequential heterodonty is lower (P_HMS_=0.0755), and dignathic heterodonty significantly higher (P_HDG_=0.0058) in Squalomorphii.Overall, these findings do not generally support strong superorder-level biases that might facilitate, or canalize, dental variation within individuals.

**Figure 2.**
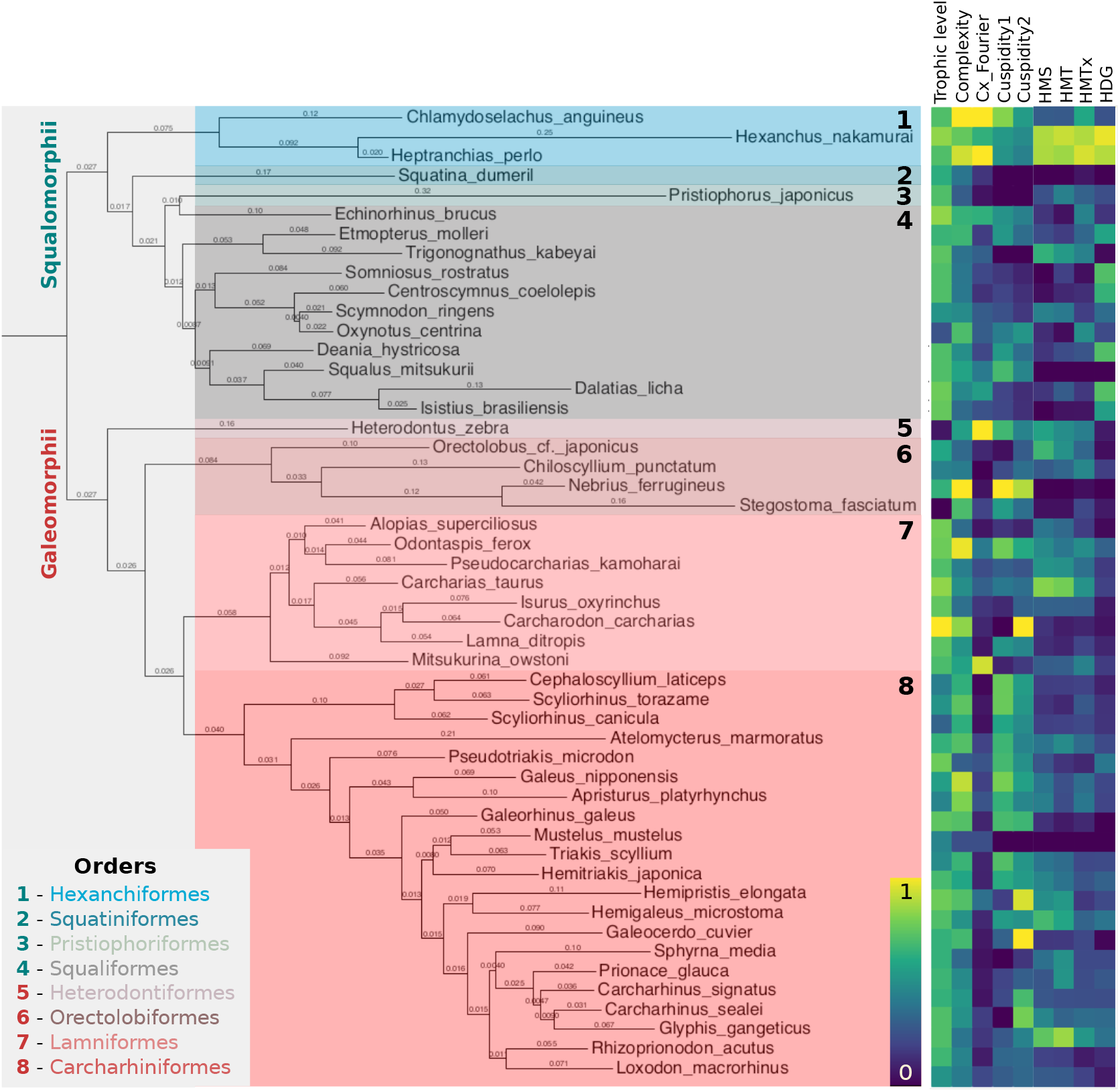
Heterodonty is widespread across all shark clades. We selected 51 species across the entire *Selachimorpha*, representing most of the extant shark diversity. Displayed branch lengths are proportional to genetic distance (see Methods) and taxonomic orders are distinguished by background (and font) colour. The adjacent heatmap shows species-wise measures of trophic level, tooth-level complexity (average of different measures, see FIG.11), Fourier-based tooth-level complexity, cuspidities (1:coarse, 2:fine), and heterodonty measures, each normalized by the respective global minima and maxima: HMS-sequential monognathic heterodonty, HMT-total monognathic heterodonty, HMTx-maximal heterodonty between any two teeth of the same jaw, HDG-dignathic heterodonty.

### 2. Heterodonties are correlated

It is conceivable that some species may show a high degree of between-jaws tooth variation without exhibiting high variation between adjacent teeth, and *vice versa*. However, we find strong correlations between monognathic and dignathic heterodonties (HMT∼HMS: R=0.9197, HDG∼HMT: R=0.4236, HDG∼HMS: R=0.3062), suggesting that variation within and between jaws is not independent (FIG.3); the main outliers being squalean, which, on average, exhibit low monognathic but high dignathic levels of heterodonty (FIG.9A). The highest levels of both monognathic and dignathic heterodonty are found within Hexanchidae featuring highly specialized dentitions, whereas *Mustelus, Squatina, Nebrius* and *Squalus* occupy the opposite end of the distribution. Strikingly, the latter genera exhibit very different tooth morphologies, ranging from plaque-like (*M*.*mustelus*), unicuspid (*S*.*dumeril*), to asymmetrically bent (*S*.*mitsukurii*) and complex, multicuspid, teeth (*N*.*ferrugineus*), indicating that there may be no trivial correlation between tooth-level complexity and dentition-level complexity. This is quantified by low-to-medium correlations between different complexity and heterodonty measures (HMS∼Complexity: R=0.196, HMT∼Complexity: R=0.1275, HDG∼Complexity: R=-0.063). Overall, most galeomorphs show a more gradual heterodonty pattern than squalomorphs, as measured by the ratio of the maximal shape difference and HMS between any two teeth (FIG.9B).

**Figure 3.**
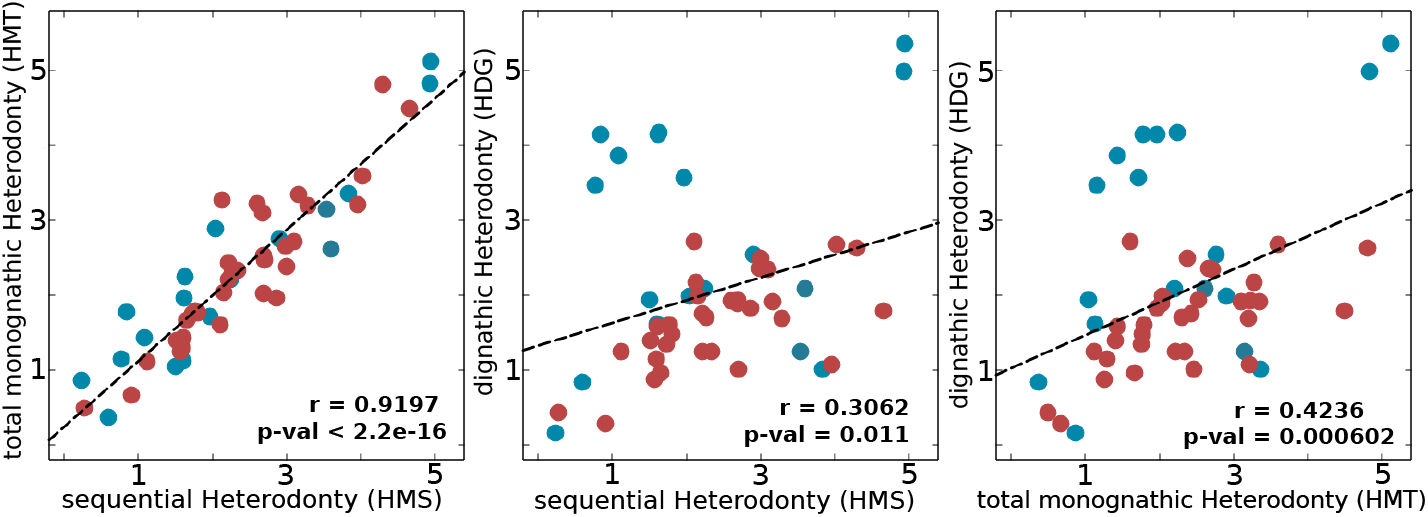
Heterodonty measures are correlated. Sequential (HMS) and total (HMT) monognathic heterodonties, as well as dignathic heterodonty (HDG) are plotted against each other, revealing positive correlations. Colors encode phylogenetic clades (blue: *Squalomorphii*, red: *Galeomorphii*). P-values are based on Pearson’s correlation test.

### 3. Patterns of heterodonty and dental complexity across phylogenetic distances

The widespread occurrence of heterodonty and tooth-level complexity (cf.FIG.2) across the entire shark phylogeny suggests a repeated evolution of these features. In order to test how dynamically these features vary at different phylogenetic scales, we correlated genetic distances with differences in dental morphological descriptors. This analysis reveals that between genetically close or moderately distant species, there is no significant positive correlation between genetic and heterodonty increases (FIG.4, see also FIG.15). Only the genetically most distant species exhibit significantly higher heterodonty differences. This finding implies that heterodonty can change relatively unconstrainedly, suggesting substantial evolvability. Conversely, differences in tooth-level complexity increase significantly between very closely and intermediately related species. However, this does not apply to every specific measure of tooth-level complexity individually, as differences in cuspidity are uncorrelated with genetic distance (FIG.15). Overall, our findings suggest that some tooth-level complexity measures might serve as a moderately better proxy for relatedness than heterodonty.

**Figure 4.**
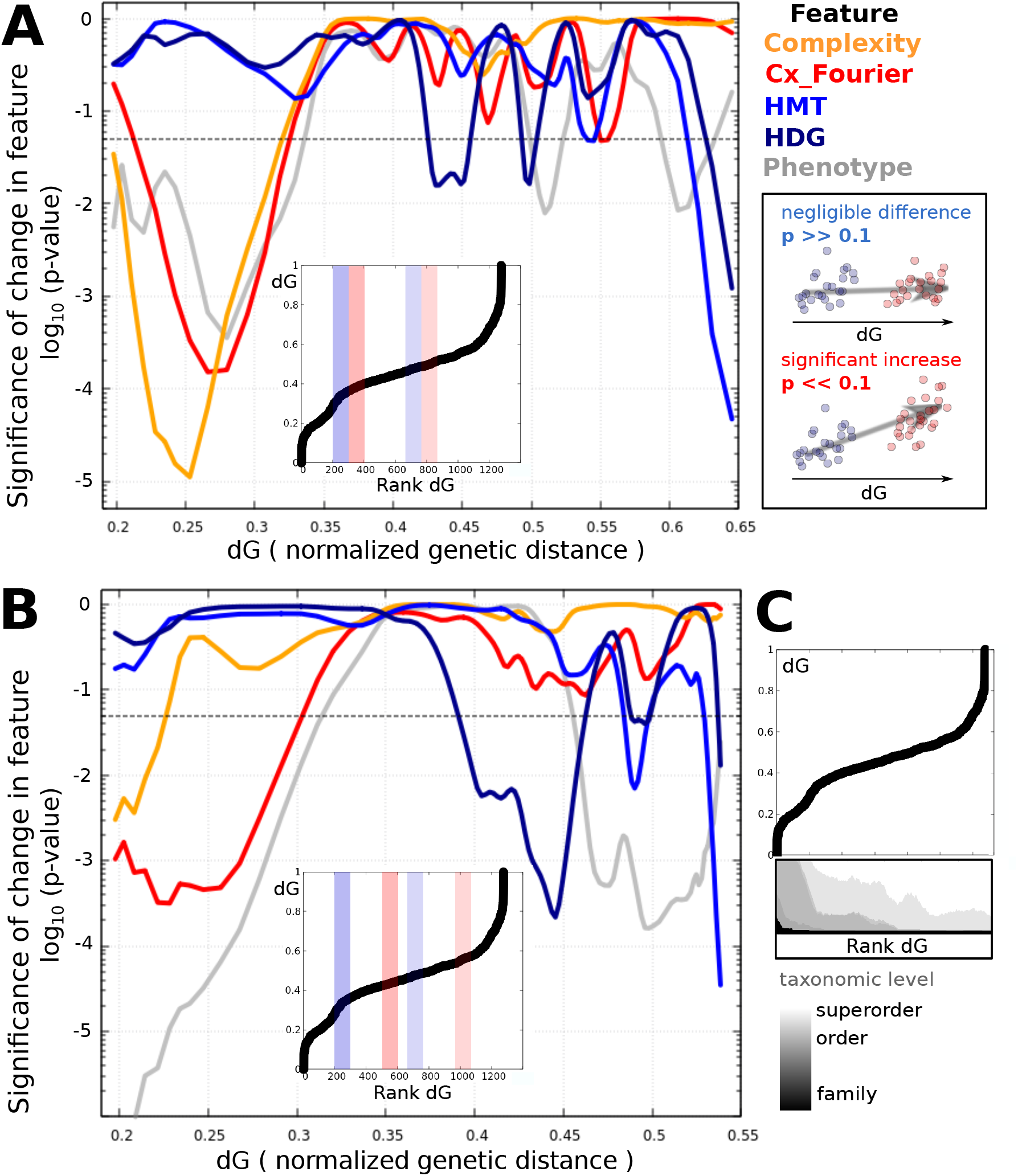
Differences in heterodonty show no significant increase with genetic distance for low-to intermediate taxonomic levels, unlike tooth-level complexity. We ordered all pairs of species by normalized genetic distance (dG) and calculated the p-values (one-sided Wilcoxon test) for significance of difference of overall tooth-level complexity (orange), Fourier-based complexity (red), total monognathic heterodonty (HMT, blue), dignathic heterodonty (HDG, dark blue) and total phenotypic distance (grey) between two subsets of 100 species pairs each. Total phenotypic distance is based on position-wise tooth shape comparisons between different species. Subsets were defined as containing the nth to the n+100th species pair ordered by dG, for sliding (incrementally increasing) n. The two subsets were **A)** subsequent or **B)** 200 ranks apart, in order to account for different scales of comparison. Here, the lines connecting the p-values are Bezier-smoothened and plotted against dG of the highest member of the lower set of species pairs. A dotted line marks the 0.05-level of statistical significance. Inlets show the relationship between dG and ordered ranks and examples of two pairs of subsets (higher:red, lower:blue). For illustrative purposes, schematic examples of two pairs of set with low (red) and high (blue) p-values are displayed besides. **C)** For orientation, we display the taxonomic compositions of the ordered species pair sets, with black representing the portion of pairs from the same family (only few), light grey representing pairs from the same superorder, and white pairs stemming from different superorders, with intermediate shades of grey referring to intermediate taxonomic levels.

Despite the absence of a substantial mid-range phylogenetic signal for specific heterodonty measures, we tested whether combinations of these measures differ between main shark clades. Using canonical correlation analysis (CCA), we are able to separate Squalomorphii and Galeomorphii, the two shark superorders (FIG.5). While including only morphological features into the CCA allows to separate the Squalomorphii from 80% of Galeomorphii along the first canonical axis (FIG.5A), adding some key ecological features leads to a complete separation of the two superorders (FIG.5B). Interestingly, the ratio between monognathic and dignathic heterodonty, and graduality of heterodont change, appear to be better superorder separators than each heterodonty measure in isolation (FIG.5C). Consistent with the higher correlation between differences in dental complexity and genetic distance, we find that certain tooth-level complexity measures, such as fine cuspidity (and the ratio between fine and coarse cuspidity) and Fourier-based complexity, show a significant clade-specific range of values. Finally, our study also reveals significant clade-specific differences in ecological parameters, especially depth, suggesting that the identified heterodonty and tooth complexity patterns may at least partly represent patterns of adaptive morphological changes.

**Figure 5.**
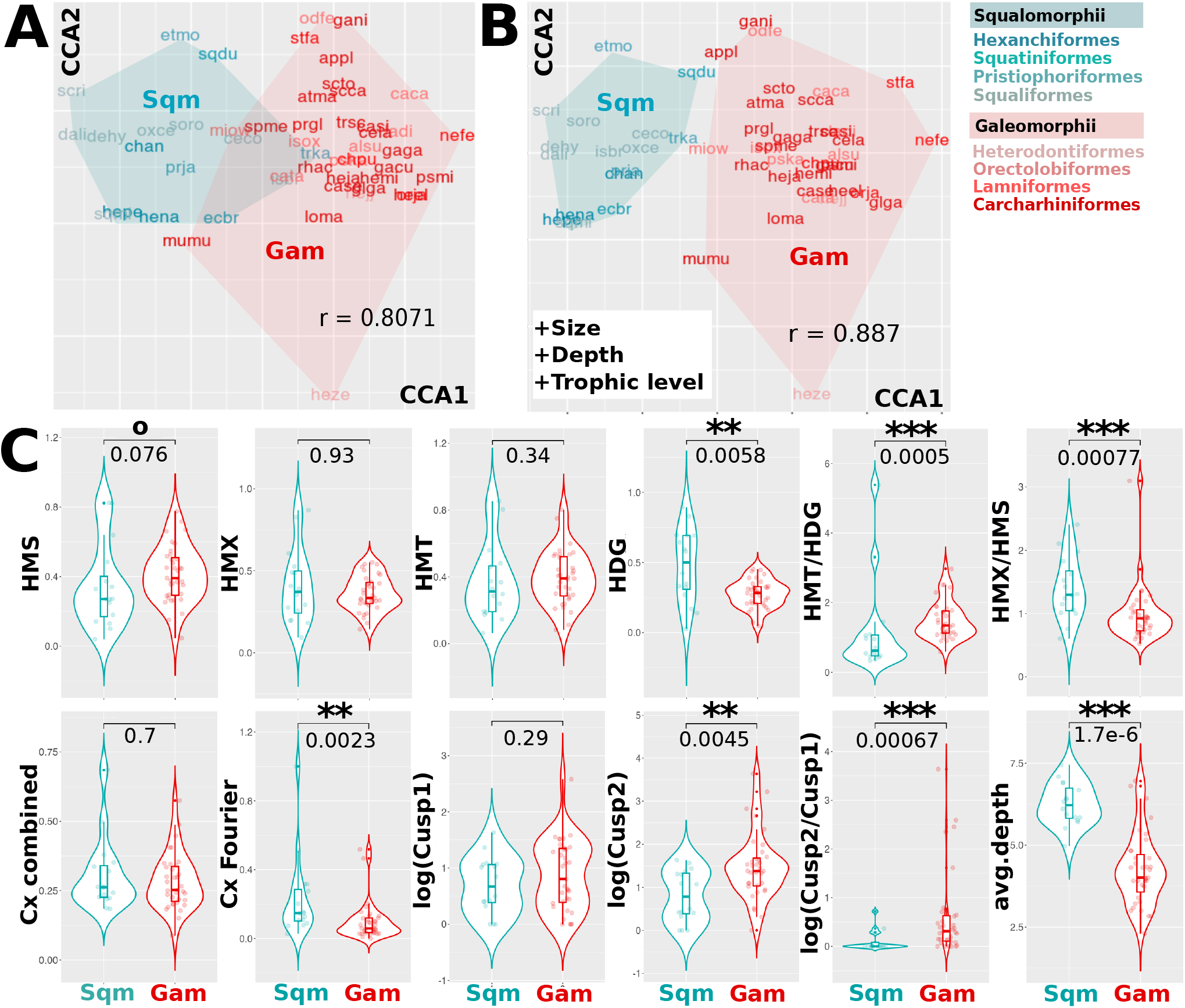
Heterodonty and tooth-level complexity measures separate shark superorders. **A**) Canonical correlation analysis (CCA) reveals combinations of heterodonty and tooth-level complexity measures that are specific for the two superorders, squalean (Sqm, turquoise) and galean (Gam, red) sharks. **B**) The two main clades are separated more clearly if ecological traits are included into the canonical analysis. The colors of the displayed species acronyms correspond to the respective orders, as displayed besides. **C**) Violin plots contrast specific features and feature combinations in Squalomorphii and Galeomorphii, with p-values plotted above (Wilcoxon test). Monognathic and dignathic heterodonty, the ratio between the two, Fourier-based tooth-level complexity (Cx_Fourier), and heterodonty and cusp ratios, as well as depth, show significant differences between the two clades. Cx_combined is the sum of all tooth-level complexity measures, Cusp1 and Cusp2 are coarse and fine cuspidity, respectively. Significance: 0.1>p>0.05: °, 0.05>p>0.01: *, 0.01>p>0.001 : **, 0.001<p : ***.

### 4. Correlations between ecological traits and heterodonty

Since tooth shapes and their arrangement within dentitions are expected to be fine-tuned towards specific niches, we evaluated correlations with ecological trait proxies. Specifically, we collected information on prey categories coarsely associated with different trophic guilds or feeding strategies (cephalopods, other molluscs, crustaceans, unspecified worm-shaped animals, non-osteichthyan vertebrates, trophic width, trophic level), habitat categories (shore, shelf, deep sea, open ocean, reefs, benthos and marine nectos, depth and depth range), and body size. We used both linear correlation models and canonical variate correlations between dental measures and ecological features (FIG.6). Although partially overlapping, these two approaches yield distinct profiles, owing to different methodologies. Dignathic heterodonty shows some correlations with habitat classes, whereas monognathic heterodonty measures emerge as important diagnostic predictors in the canonical analysis paradigm. Intriguingly, we find stronger correlations between dental features and habitat than trophic categories. An inverse correlation pattern separates residents of shallow and open-sea habitats, with species inhabiting the deep sea presenting another pattern of correlation with heterodonty and tooth complexity traits (FIG.18). Across trophic guilds, differences appear between nectic (vertebrates and cephalopods) versus bottom-dwelling prey classes (crustaceans, non-cephalopod molluscs and diverse small invertebrates). Finally, coarse cuspidity is distinctive for the difference between bottom-vs.-water column feeding strategies, actively hunted prey, and body size, while fine cuspidity is informative about size, and deep-vs.-shallow habitat distinction. Taken together, we observe distinct correlation patterns between dental descriptors and ecological traits.

**Figure 6.**
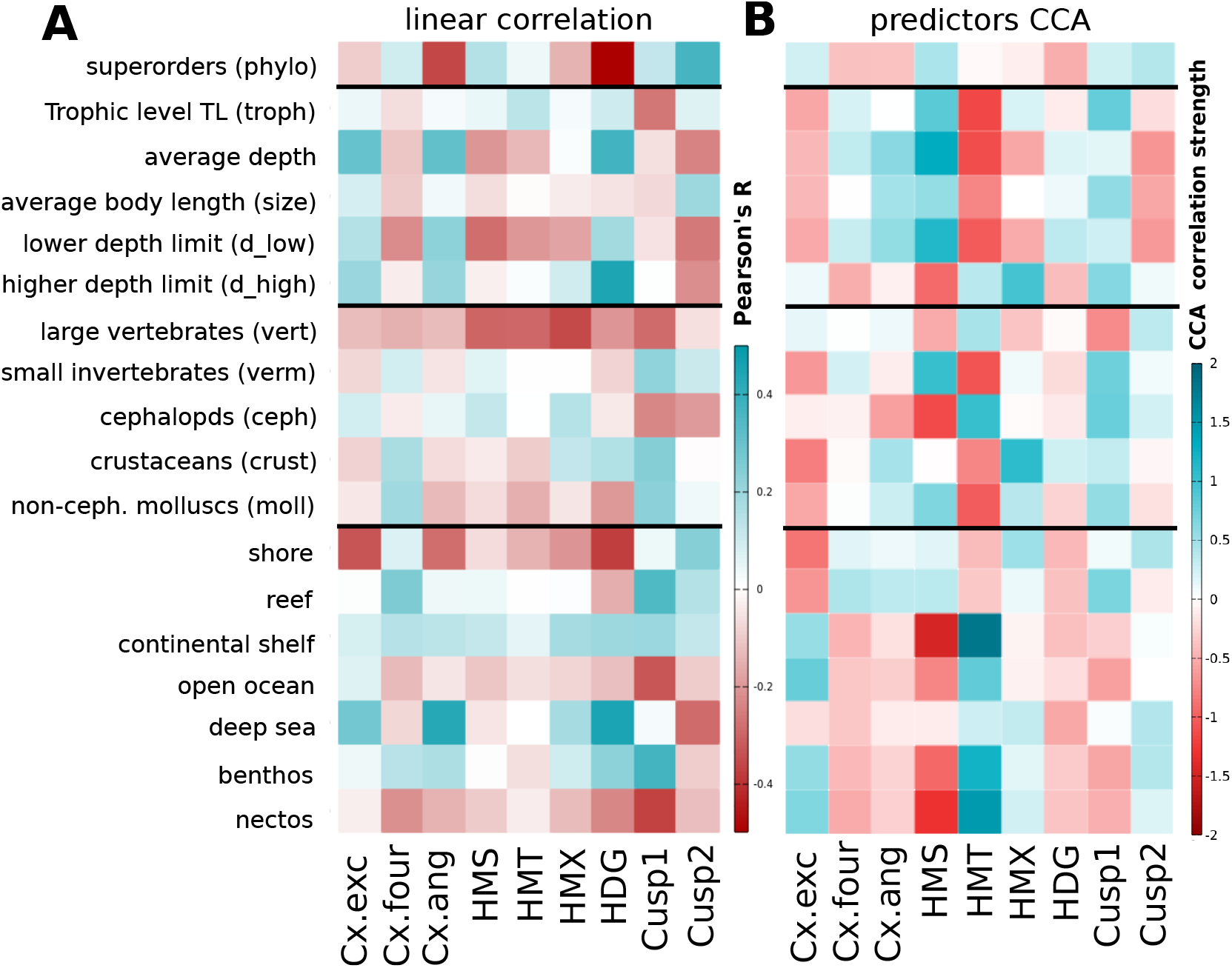
Overview of correlations between heterodonty/complexity measures and ecological traits. **A)** Different heterodonty measures, and measures of tooth-level complexity, show specific correlations with ecological features, such as body size, depth, prey guilds and habitats. Intensities of red and turquoise display Pearson’s correlation coefficient R. **B)** Analogously, canonical correlation analysis reveals that monognathic heterodonties are central in statistically separating different habitats and trophic traits. Red and blue hues indicate correlation strengths of CCA1 for linear combinations of the predictors, i.e. tooth level complexity and heterodonty measures (red: negative correlations; blue: positive correlations). Cx.exec: comprises tooth-level complexity measures based on excentricity (OCR, OAR, OIR), Cx.four: Fourier-based complexity (DFS) and Cx.ang: angle-based complexity measures (ANS, ASC, AND, OPC).

### 5. Two contrasting strategies emerge

In addition to phylogenetic and ecological groups, we find that shark species diverge along two disparate directions when plotting monognathic heterodonty against Fourier complexity (FIG.7A). While the first group (G1) shows high Fourier tooth-level complexity, but low monognathic complexity, the second one (G2) presents the reverse pattern. Teeth in G1 tend to be smoother, more obtuse and asymmetric relative to G2. Significant differences emerge when comparing the ratios between (a) coarser and finer cusp numbers, (b) different heterodonty measures, and (c) outline-vs.angle-based tooth similarity or complexity measures (FIG.7C). Leveraging combinations of these measures reveals a specific shape pattern, with G1 featuring asymmetric, compact teeth that vary little within jaws, and G2 featuring more excentric or triangular teeth that are part of morphologically heterogeneous gradually changing dentitions. Interestingly, trophic differences between the groups are not salient. However, species belonging to G1 tend to inhabit deeper regions, while G2 species are found closer to the surface, including both proximal (shores) and distal (open ocean) environments (FIG.7B,C).

**Figure 7.**
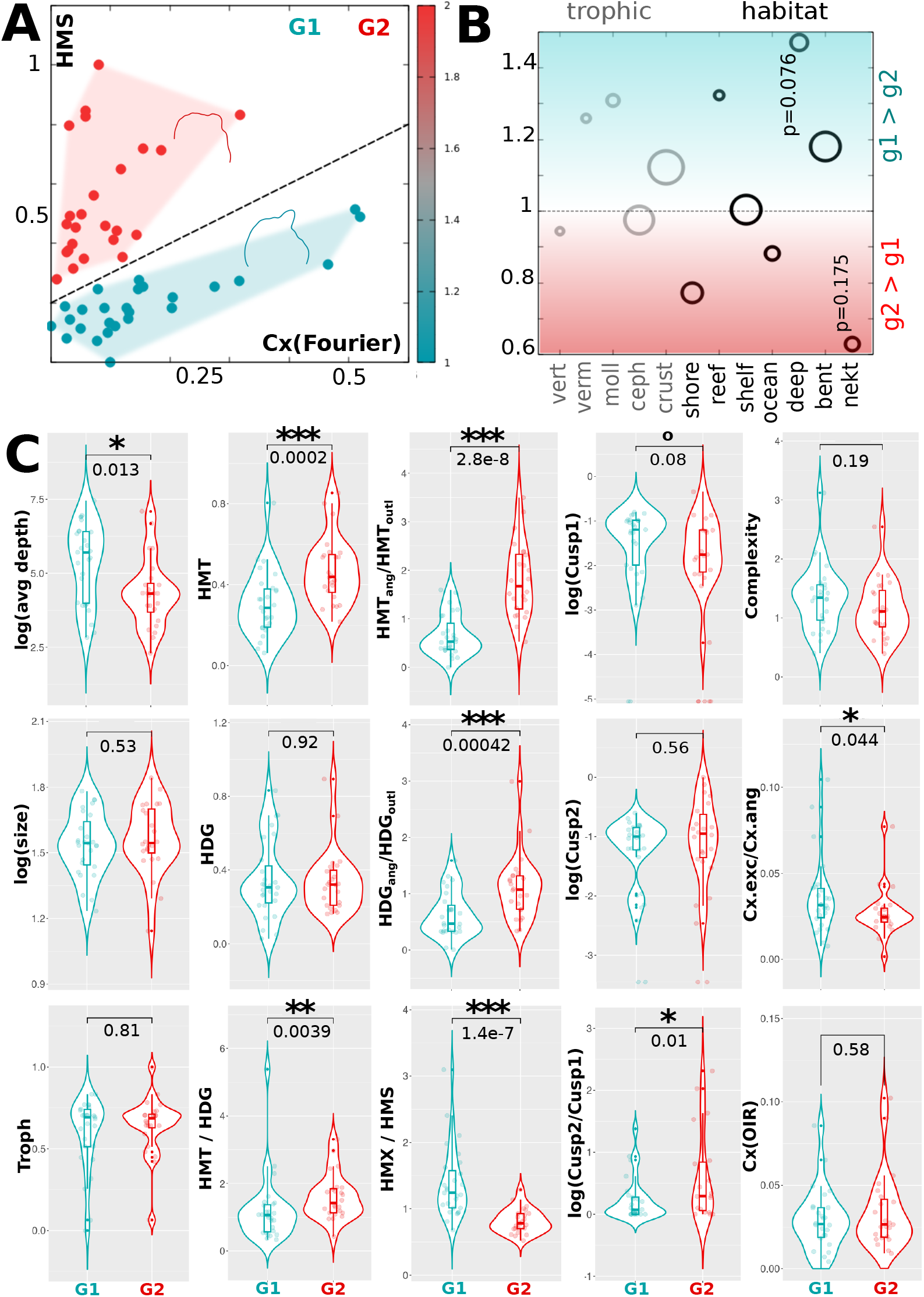
Combinations of heterodonty and tooth-level complexity measures reveal two distinct strategies. **A)** Many shark species show, roughly, either high Fourier-based complexity and low monognathic heterodonty, or *vice versa*, but rarely exhibit high values for both. Teal marks the former (G1), red the latter group (G2). Shown outlines display, respectively, mean tooth shapes for both groups. **B)** Group-wise enrichments of trophic and habitat features, each ring presents the ratio between the respective percentages of species belonging to G1 divided by the ones belonging to G2 and the expected unbiased ratio. Ring size reflects the number of species per ecological category. p-values (Student’s t-test) are annotated for the most significant differences. **C)** Violin plots visualizing further group-specific characteristics with p-values (Wilcoxon test). Notably, the groups show divergent ratios between heterodonty measures based on outlines (X_outl_) and outline angles (X_ang_), corresponding to the heterodonty measures EMD, HED and SAO vs. OAD and ADD (cf.Methods). Cx.exc comprises complexity measures based on excentricity (OCR, OAR, OIR), Cx.ang (ANS, ASC, AND, OPC) measures based on angle complexity. Cx(OIR) is the minimal ratio between the areas of inscribed and escribed circles. Significance: 0.1>p>0.05: °, 0.05>p>0.01: *, 0.01>p>0.001 : **, 0.001<p : ***.

### 6. Monognathic heterodonty is the most relevant predictor for ecological traits

As measures such as cuspidity quantify complexity on a single-tooth level (the unit), heterodonty refers to complexity resulting from the higher-order composition of these units. Based on this realization, our dataset is suitable to analyze to which extent complexity on each of those different scales matters ecologically. As a proxy for relevance, we compared correlation strengths of CCA1 coefficients (cf.FIG.6), comprising different heterodonty and tooth-level shape descriptors used to separate sharks by ecological traits or phylogenetic group, respectively. Globally, monognathic heterodonty shows particularly high correlation strengths, while fine cuspidity, in particular, yields significantly lower values (FIG.8). In detail, we also observe high correlations between food-related canonical variates and coarse cuspidity as well as dignathic heterodonty and depth, whereas body size is the trait most significantly correlated with fine cuspidity. Overall, we conclude that intermediate complexity levels (namely monognathic heterodonty) are the best predictors of most ecological traits.

**Figure 8.**
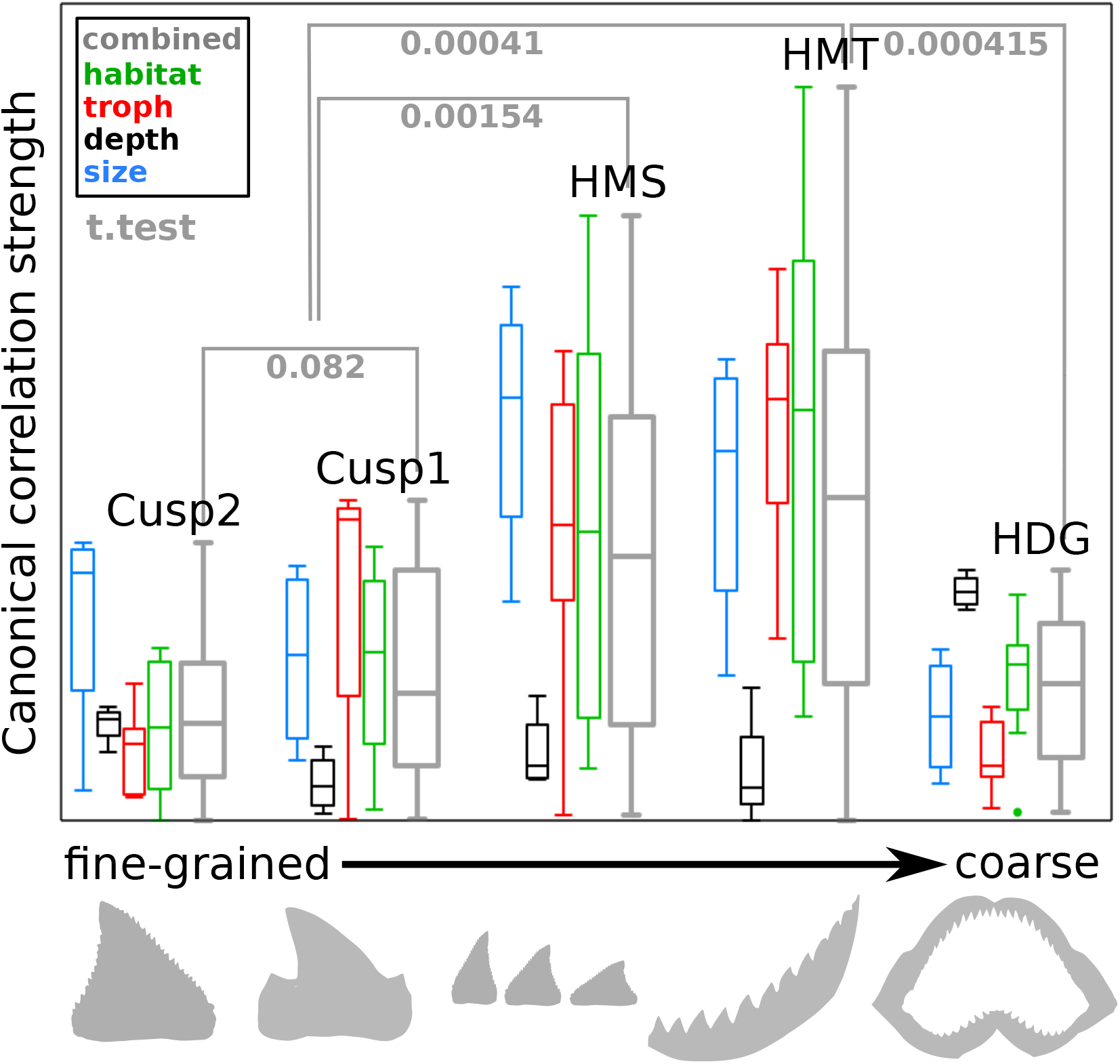
Ecological relevance of dental shape descriptors varies across resolution levels. Box plots summarizing canonical correlation strengths between different cuspidity and heterodonty measures, and heterogeneous ecological traits. Correlation strengths were found highest for monognathic heterodonty, while fine cuspidity yielded the lowest average correlation. As those measures represent, roughly, morphological trait differences (or complexity) on different relative resolution levels from fine cusps to differences between jaws, they are plotted in ascending order from the finest to coarsest scale. Canonical correlation strength can be used as proxy for average relevance, suggesting size scale-dependent differences in ecological trait importance (cf.FIG.6). Colors separate features, while the grey boxes contain the combination of all traits. Displayed p-values were calculated using a Student’s t-test. Underlying grey shapes are included for illustrative guidance.

**Figure 9.**
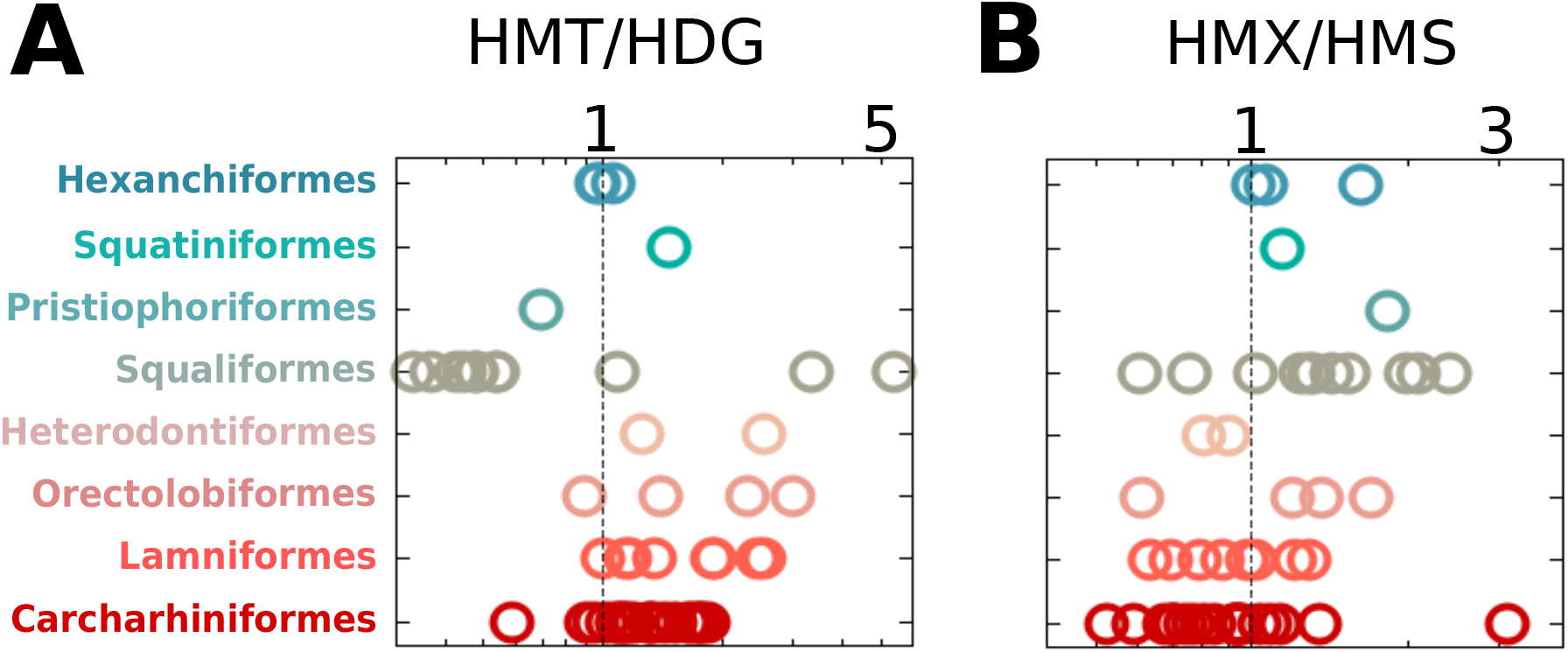
Heterodonty ratios exhibit differences across shark phylogeny. Ratios between monognathic and dignathic heterodonty (HMT/HDG) and between maximal monognathic heterodonty and sequential monognathic heterodonty (HMX/HMS, a proxy for graduality, with lower values indicating higher graduality), show different values between shark orders. Here, shark orders are lined up along the y-axis and are marked by colours; the x-axes denote heterodonty ratios using log scales. Notably, Squaliformes exhibit high dignatic and low monognathic heterodonty, while galean sharks show overall more gradual shape change than squalean sharks.

**Figure 10.**
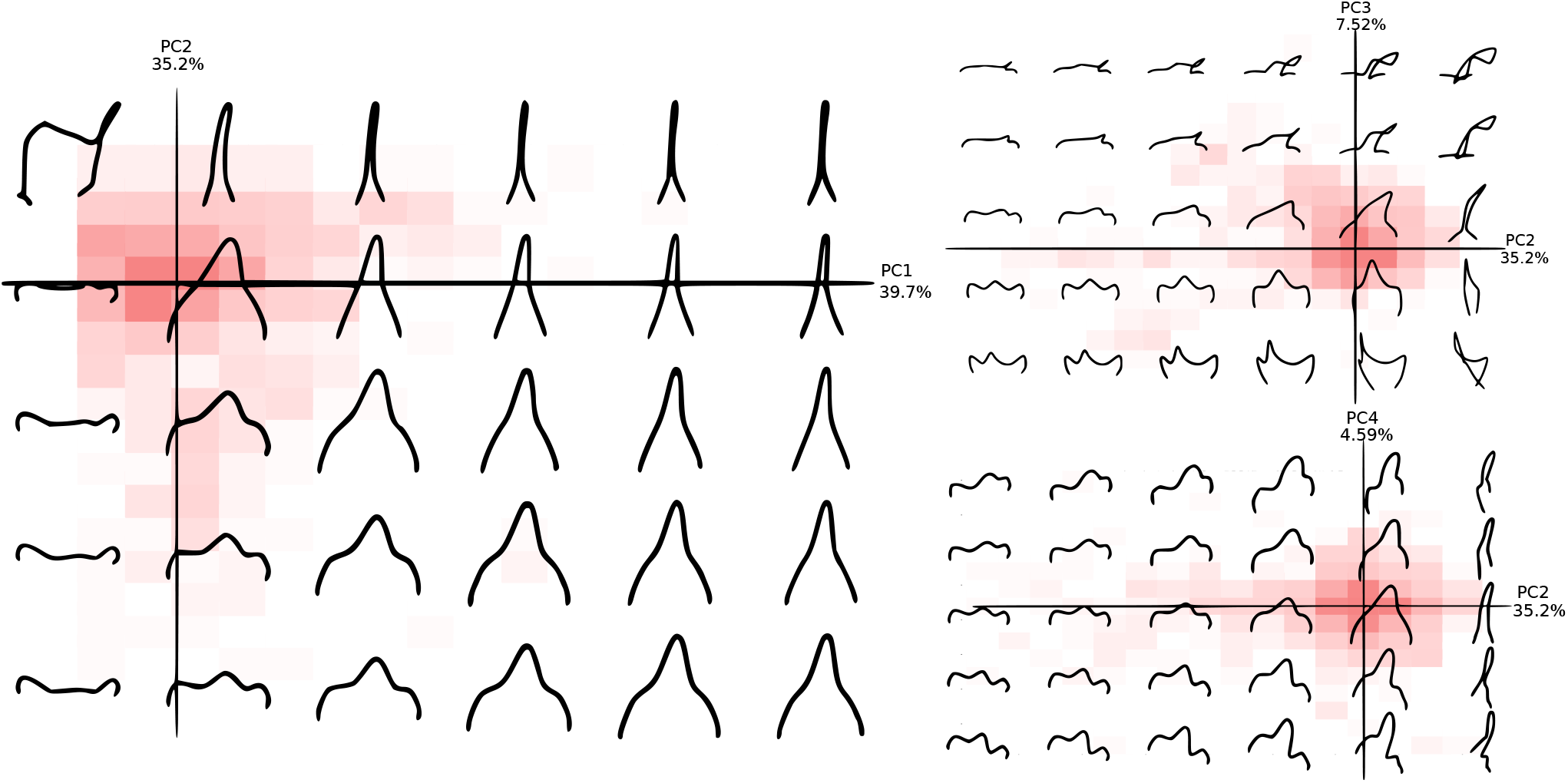
Single tooth discrete Fourier PCA. We compared individual tooth shapes taking advantage of the discrete cosine Fourier analysis. Here, we show calculated theoretical outlines for combinations of the first four PCs superimposed over binned densities of their respective occupancies across all shark teeth used in the analysis. Higher densities correspond to darker 2D bin colour.

**Figure 11.**
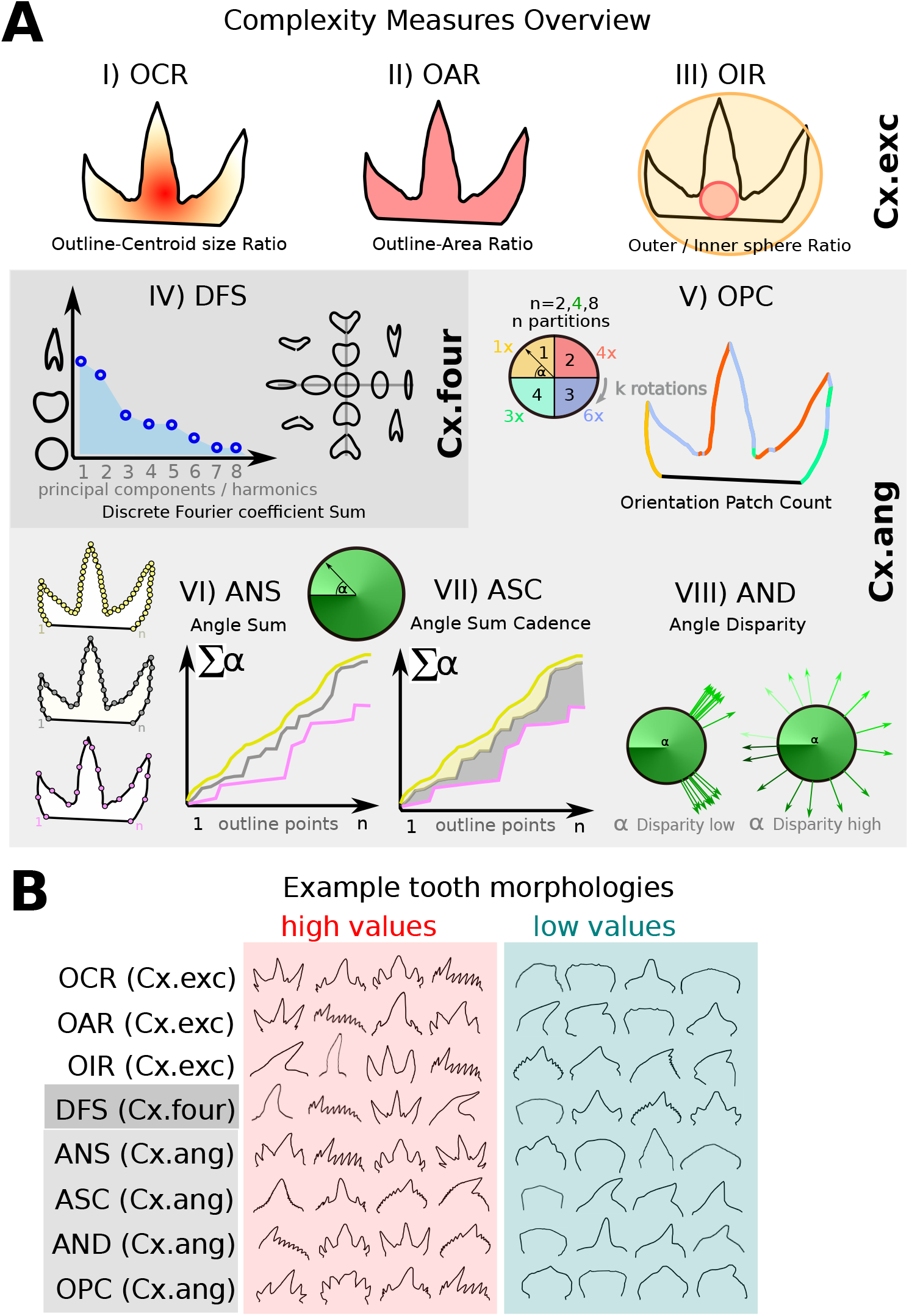
Overview of tooth-level complexity measures. **A)** Schematics illustrating the different methods. Different background shades separate different groups of measures that tend to be correlated FIG.12: Cx.exc, Cx.four, Cx.ang. i) OCR: ratio between outline length and centroid size. ii) OAR: ratio between outline length and total area. For these methods, high complexity is associated with excentric shapes with large cusps or protrusions. iii) OIR: ratio between area of incircle and excircle. Excentric and gracile shapes yield high complexities. iv) DFS: describing more complex and excentric shapes requires more Fourier harmonics. In the example morphospace, more central shapes featuring smaller coefficient values, are more similar to a circle. Thus, this measure does not attribute high values to reiterative patterns, such as equal cusps. v) OPC: outline angles are discretized according to an absolute coordinate system with different partition numbers. Frequent change of discrete angle numbers along the outline is associated with complexity of outline features. vi) ANS: sum of angles between outline points for a set of resolutions (here exemplified with different colors), attributing complexity to outline feature density. vii) ASC: this measure quantifies the difference of outline angle sums between different resolutions, i.e. it yields high values if different outline features exist on different scales. viii) AND: diversity of outline angles, i.e. feature diversity. **B)** For each complexity measure, examples of tooth shapes with high and low values are displayed.

**Figure 12.**
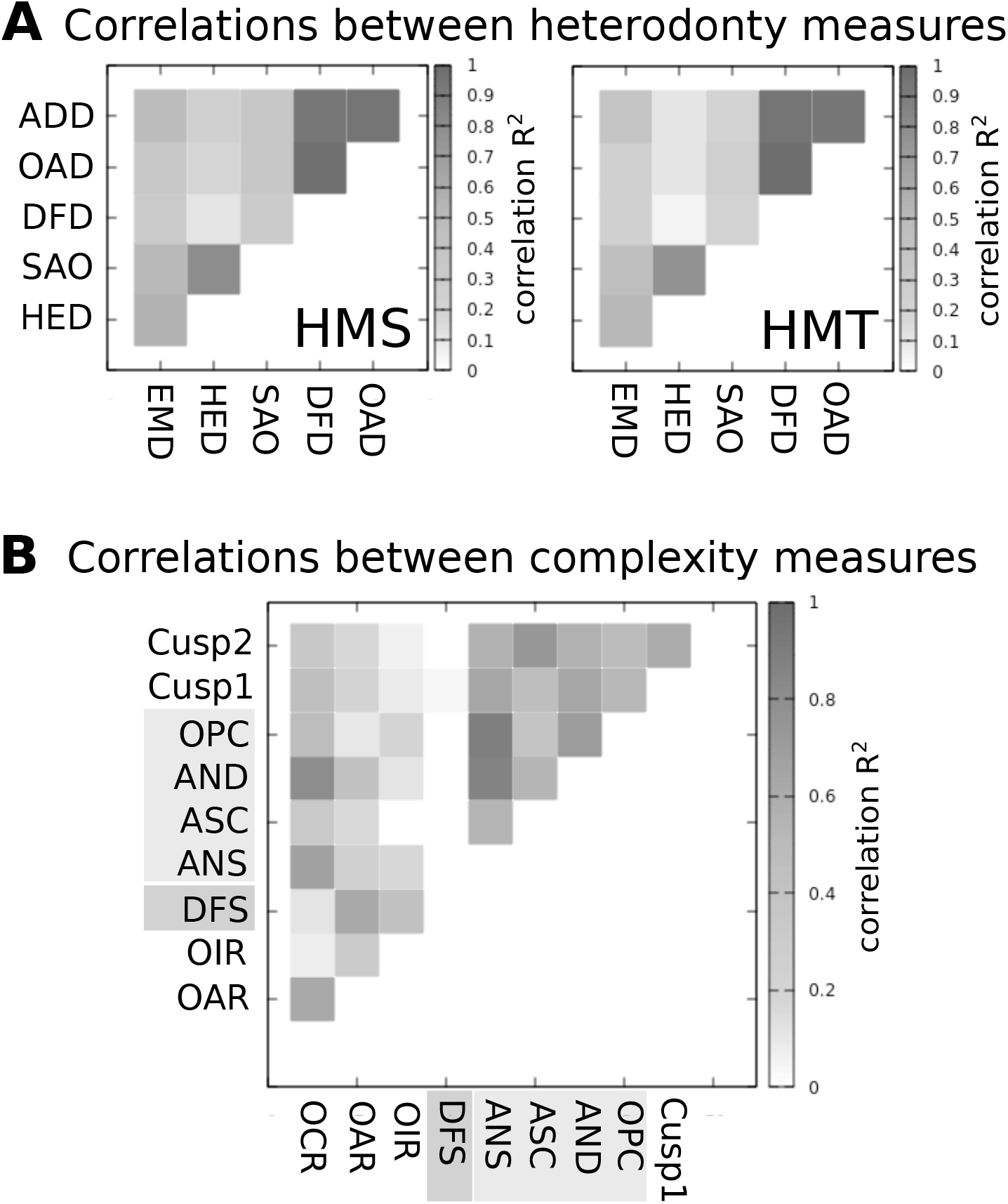
Correlations between measures. **A)** Species-wise correlations between heterodonty measures for sequential (left) and total monognathic heterodonty (right). Outline similarity measures (EMD, HED, SAO) and, especially, angle-based measures (ADD, OAD), show high internal correlations. **B)**: Pair-wise correlations for all tooth-level complexity measures, based on data for all teeth. Background shades of grey correspond to defined groups of measures (cf.FIG.11). Strong correlations exist particularly between angle-based measures, while the Fourier-based method appears only weakly correlated with the remainder, justifying the assumption of complementarity of methods. For all plots, linear correlation R values are represented by greyscale.

**Figure 13.**
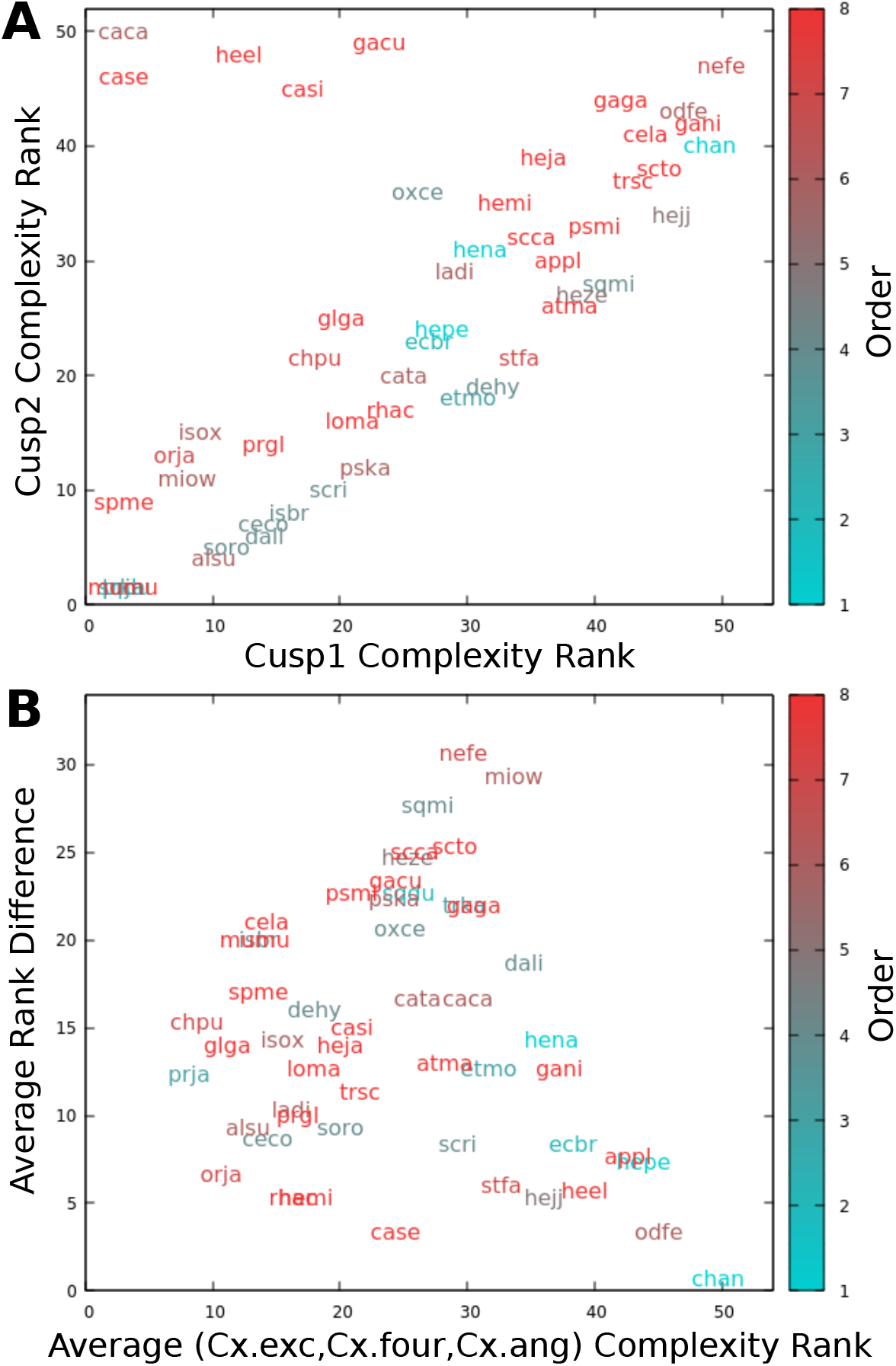
Tooth-level complexity measures are complementary. **A)** Species are ranked based on coarse (Cusp1) and fine (Cusp2) cuspidity. Colour corresponds to taxonomic order, in the same sequence as in 2. While both cuspidity ranks appear to be generally correlated, some galean species show high fine cusp numbers (typically: serrations) while maintaining an overall low coarse cuspidity. **B)** The average tooth-level complexity (based on a combination of Cx.exc, Cx.four and Cx.ang) is plotted against the average rank difference when ranks for individual complexity measures are compared, one by one. Several species show substantial differences in complexity depending on the measure applied, again suggesting complementarity of methods.

**Figure 14.**
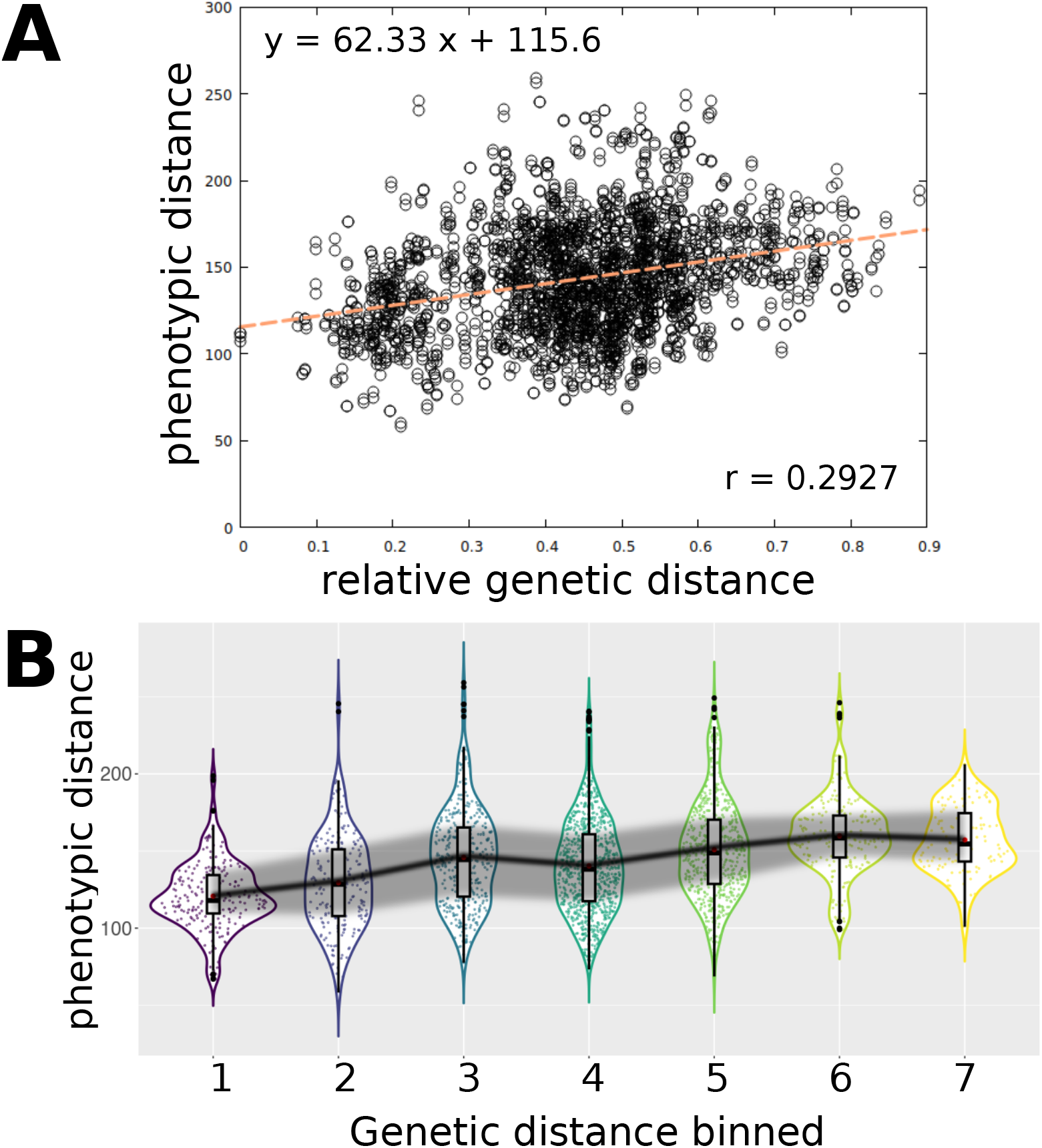
Phenotypic distance increases steadily with genetic distance. We correlate genetic distance and phenotypic distance for all pairs of shark species. Phenotypic distance is calculated as the normalized Euclidean distance of Fourier coefficients between pairs of teeth in corresponding relative positions along the jaw. **A)** correlation plot with linear regression line. **B)** Genetic distances between pairs of species were binned by deciles, collapsing the first two and highest three deciles, respectively, owing to data scarcity. Overall, a steady and almost monotonous increase of phenotypic distance with genotypic distance is observed.

**Figure 15.**
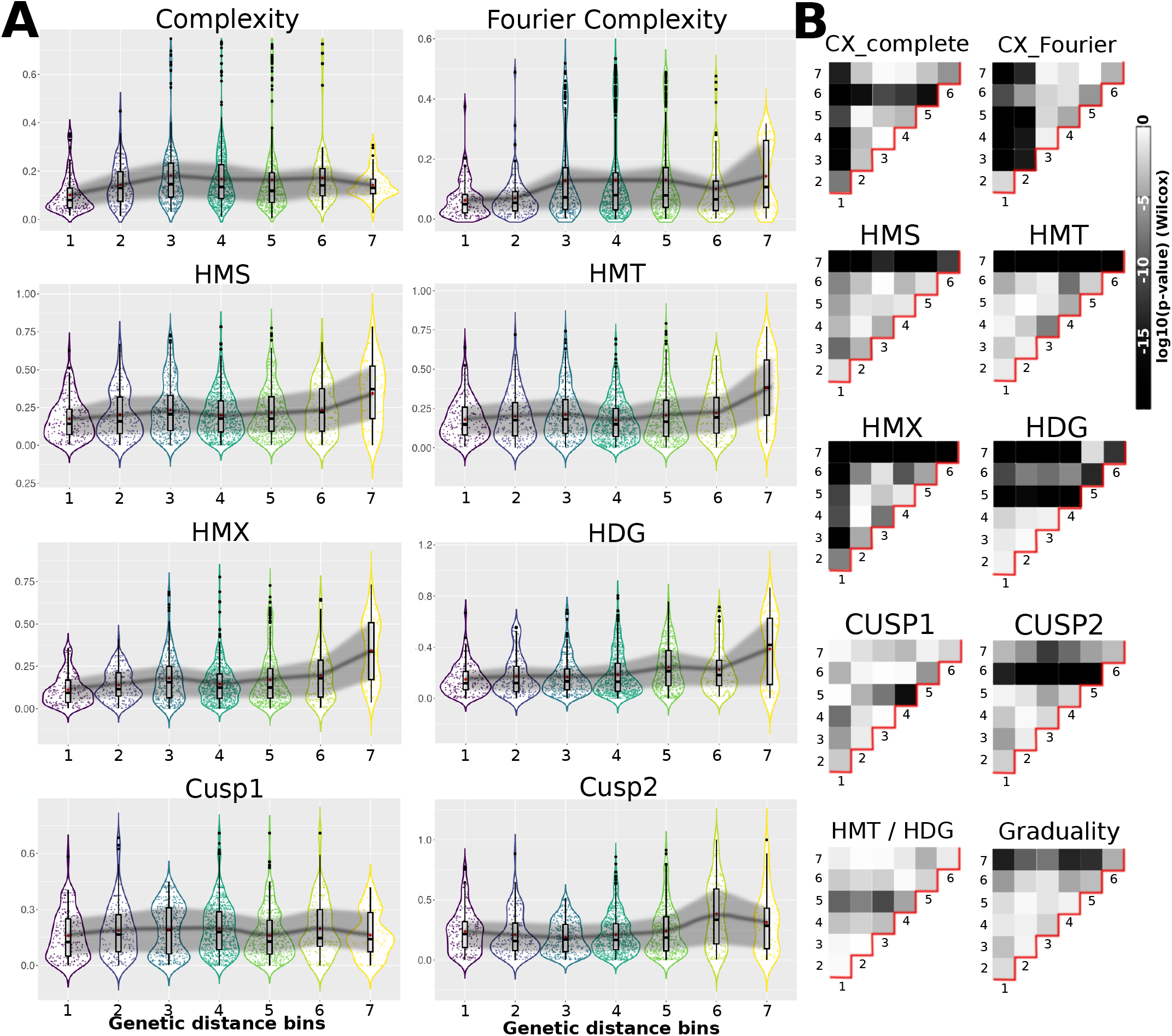
Features show different ranges of strong and weak correlations with genetic distance. **A)** Binned genetic distances between all species pairs, as in FIG.14. Corresponding differences in tooth complexity and heterodonty measures, respectively, are shown on the y-axis. For most measures excluding tooth-level complexities, no important differences are seen across most of the genetic distance range, suggesting limited depth of phylogenetic signal. The dark line connects averages per bin, with the grey shadow highlighting the range of the central 50% of all pairs per bin. B) Dissimilarity of heterodonty and tooth complexity values between bins is quantified by Wilcoxon test, with numbers on x- and y-axes corresponding to bins. values are represented by greyscale; note that they can refer to both increasing and decreasing trends.

**Figure 16.**
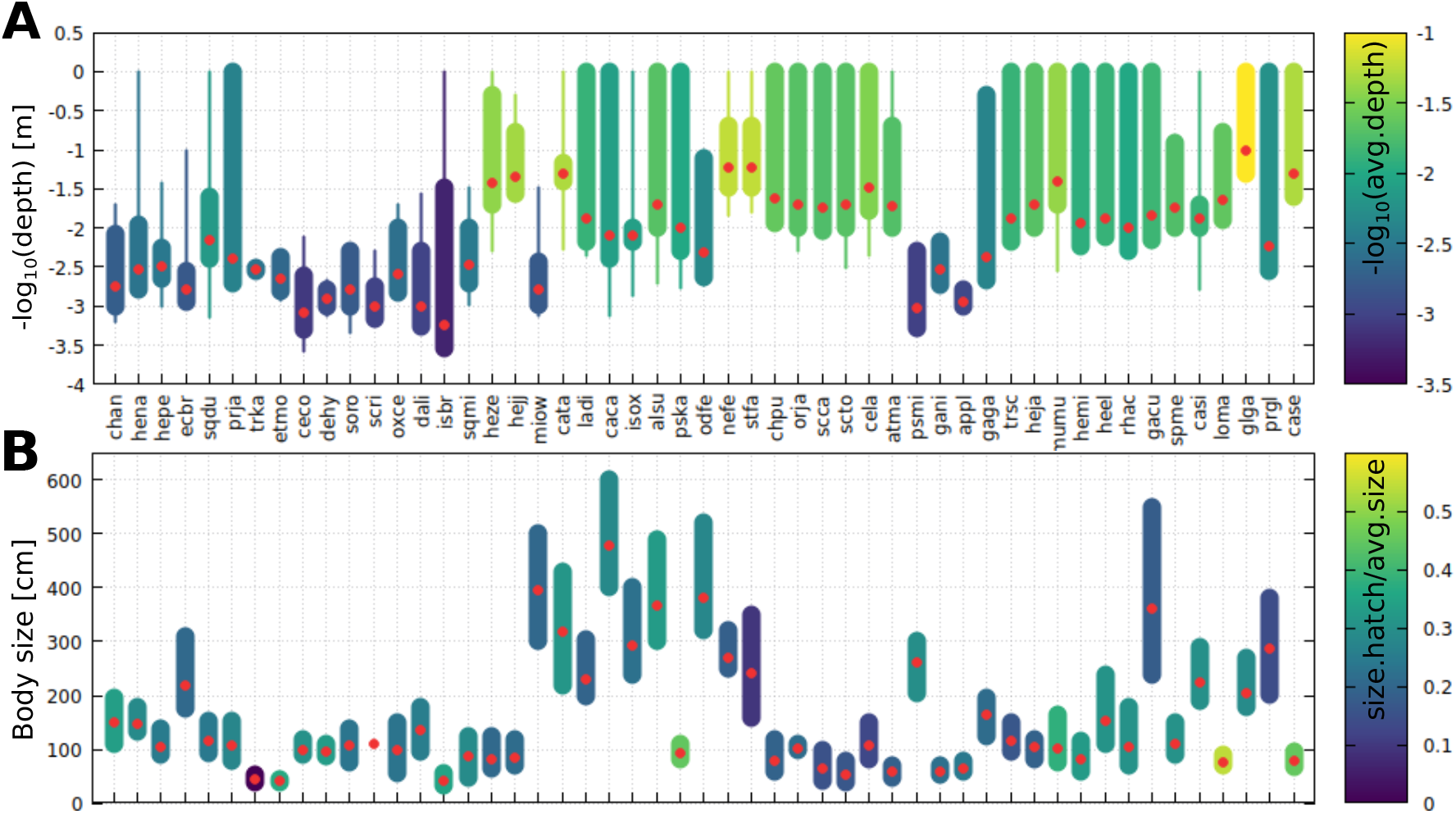
Depth and size ranges per species. This graph displays key traits linked to ecology across the scrutinized taxa. **A**) Depth: Thick bars span typical depth range per species as gathered across literature. thin candlestick ends expand to exceptional reports, red dots and the color code mark averages. **B**) Body size (length). Range of body sizes for sexually mature adults are shown. Averages of lower and upper ranges for both sexes are shown as red dots. Colors mark ratios between hatchling/newborn and average mature adult body length.

**Figure 17.**
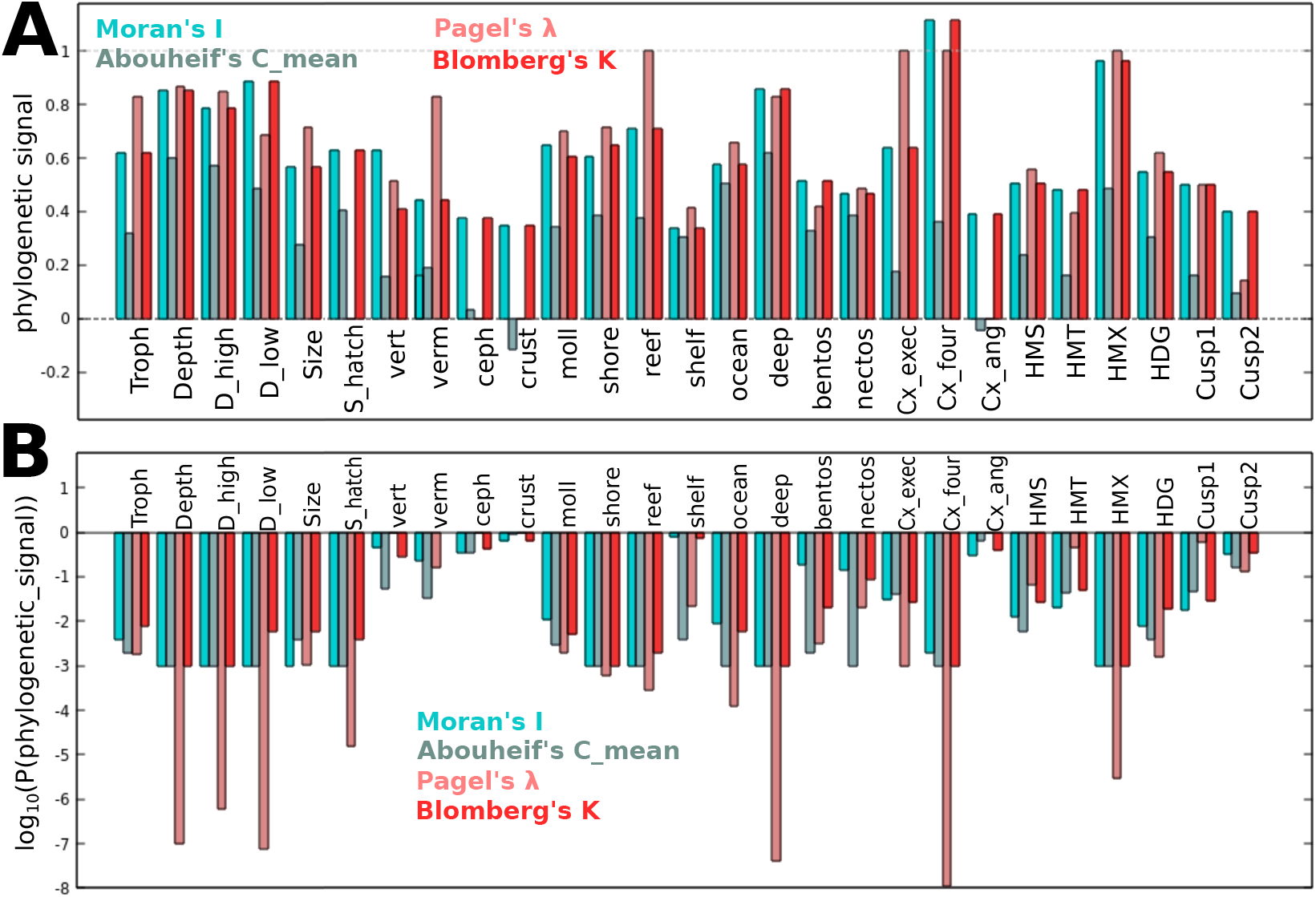
Phylogenetic signal. Using the commonly used Moran’s I, Abouheif’s C_mean, Pagel’s *λ* and Blomberg’s K, **A)** phylogenetic signal strengths and **B)** the p-values of their respective significances are plotted for ecological traits, tooth complexity, and heterodonty measures. Most trophic guilds, monognathic heterodonty measures, angle-based complexity measures, and cuspidities, only show relatively low phylogenetic signal, while some habitat traits, depth, Fourier complexity, and maximal heterodonty, exhibit strong phylogenetic signal.

**Figure 18.**
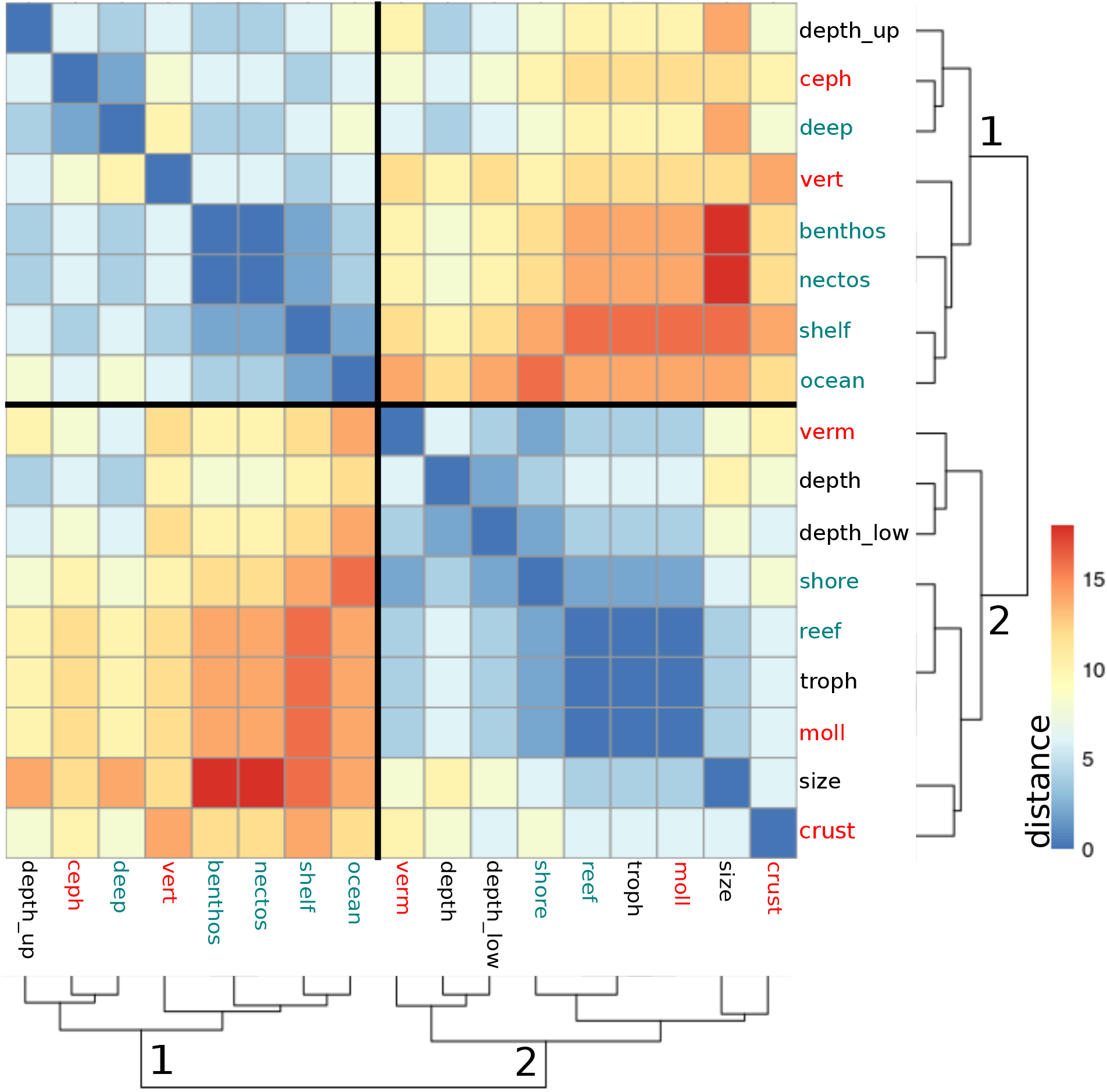
Similarity of canonical correlation profiles between ecological traits. Heatmap showing pair-wise Manhattan distances between binarized (1/-1) canonical correlation profiles for ecological traits (cf.FIG.6) with tooth complexity measures (CX1, CX2, CX3, Cusp1, Cusp2) and heterodonties (HMS, HMT, HMTX, HDG) as phenotypic coefficients. Two discernible sub-clusters emerge, pointing to discrete sets of ecological strategies that are reflected by dental patterns. Red: food guildes, turquoise: habitat specifiers.

**Figure 19.**
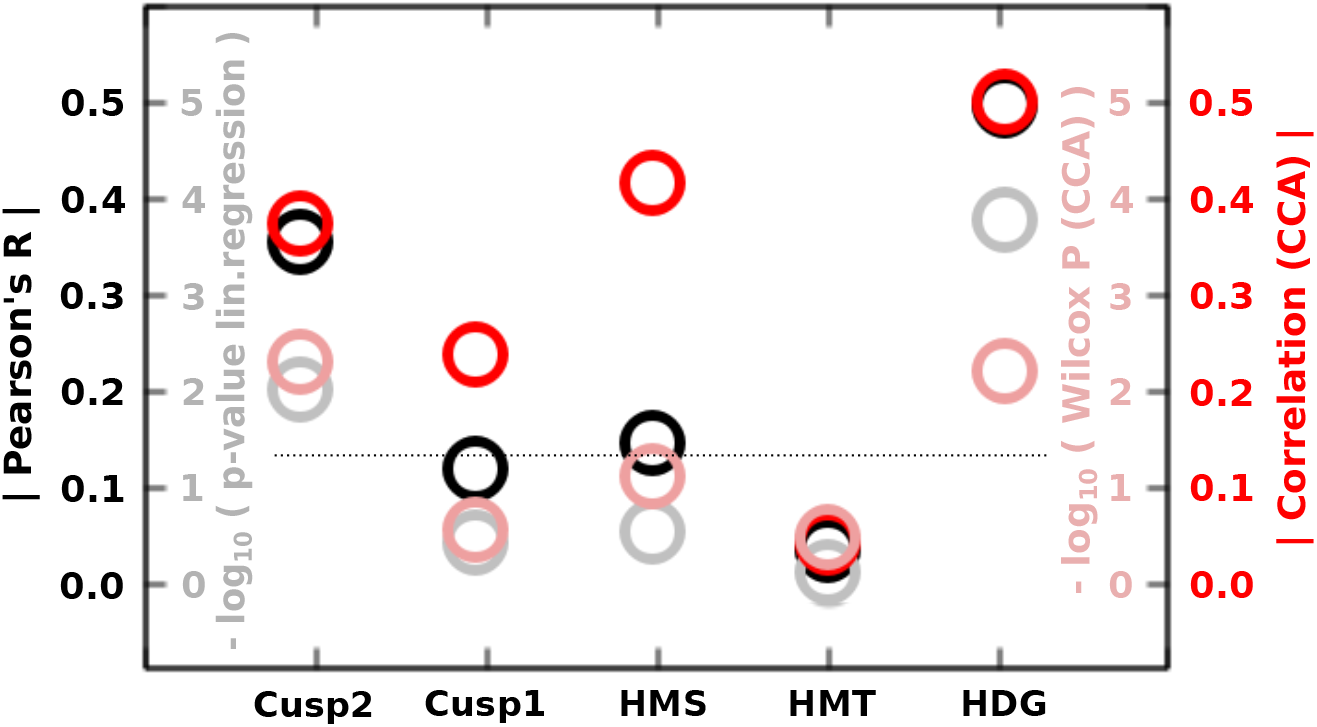
Summary: Correlations of features per resolution scale and macro-phylogeny do not reflect ecological importance. For the measures separated by resolution as in FIG.8, we show the respective correlations with macro-phylogeny (superorders) using Pearson’s correlation R, besides the corresponding p-values, CCA correlation strengths, cf. FIG.6, as well as the p-values of a Wilcoxon test, cf. FIG.5. The significance threshold of 0.05 for p-values is indicated by a dashed line. Particularly strong correlations exist for fine-grained complexity (Cusp2) as well as dignathic heterodonty (HDG), which is in contrast to the scales at which correlations with ecological proxies are most prominent, suggesting relative independence of ecological and macro-phylogenetic patterns.

**Figure 20.**
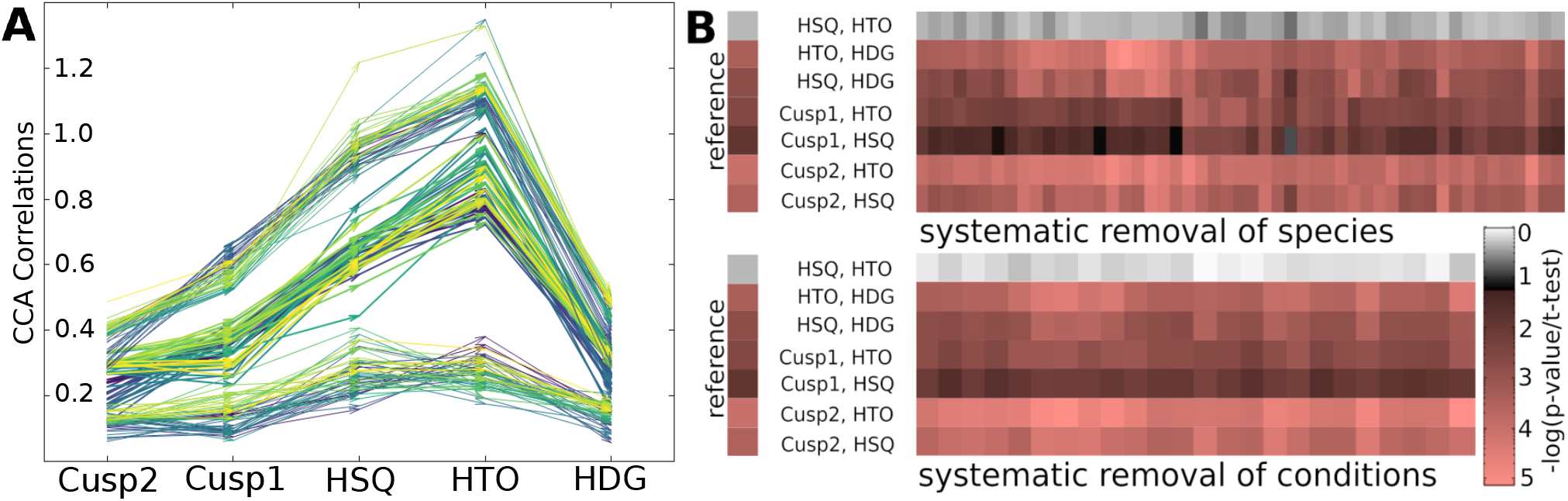
Scale-dependent ecological relevance is robust against moderate data modifications. **A)** Upon systematic removal of shark species from the data set, we recalculated CCA correlations per scale-specific complexity feature (fine, coarse cuspidity, heterodonties). For each modified data set, we plotted the three inner quartiles and connected the scales by vectors. Colors indicate which species were removed. **B)** For the same data, we remove species one-by-one (upper panel) and ecological predictors (lower panel) and plot p-values for pair-wise t-tests between scale-specific complexity measures. The color code is logarithmic and significant values (p<0.05) are highlighted in shades of reddish, with the pairs of measures indicated besides. The results displayed in FIG.8 are re-plotted as reference. It appears that the stark differences between monognathic heterodonty and the other measures shown in FIG.8 are robust against moderate alterations to the data.

## Discussion

Although separated by over 400Ma of evolution^44^, both sharks and mammals show remarkable dental variation both between and within individuals and species. We show quantitatively that, within sharks, this diversity is not restricted to specific clades but evolves dynamically. This suggests conserved developmental mechanisms capable of producing a large range of potentially adaptive tooth shapes^23,24^. Recent work has increased insight into shark odontogenesis and revealed commonalities with its mammalian counterpart^20,45^. However, important differences comprise depth of tissue embedding of tooth development, continuous generation of new teeth germs resulting in deciduous tooth series (a similar feature is present in polyphyodont mammals, too) and the local regulation of cell proliferation and apoptosis^20^. On the level of single-tooth morphologies, *in silico* approaches to mammalian and shark odontogenesis implementing some of the described differences suggest a similar capacity of generating phenotypic diversity^23,46^. However, on the level of dentitions, graduality differences between adjacent teeth are expected. This is because in mammals, Hox genes play a key role in distinguishing different tooth types along the jaw, which has not been shown yet in sharks^26,27^.

Our analysis reveals that Squalomorphii exhibit larger but less gradual variation between adjacent teeth than Galeomorphii. We suggest that the former’s lower graduality is partly due to highly heterodont dentitions in the Hexanchidae that appear to be functional and conserved throughout the proximate fossil record^47^, and to the cutting blade tooth rows widespread in Squaliformes that depend functionally on interlocked homodont teeth, thereby reducing heterodont variation^48^. Although several studies have documented gradual tooth shape variation in some galean species^29,31–34,39^, we are not aware of any previous study comprehensively quantifying heterodonty throughout the entire class. Interestingly, jaw-level differences between Squalomorphii and Galeomorphii may point to different feeding strategies that are reflected both by dentition features and jaw shape^1,40^. Strong functional adaptive pressures may also underlie increased phenotypic evolution rates in Squalomorphii^40^. While a comprehensive study analysing a single-tooth morphospace across sharks did not show clearly separated morphospace regions for orders and superorders, morphospace occupancy of Squalomorphii and Galeomorphii presented a clear bias^36^. Yet, whether development or functional constraints ultimately underlie these differences between the major clades remains to be elucidated.

Between ecological features and heterodonty, correlations emerge indicating that quantitative shape differences between teeth are functional. Specific correlations, however, may vary by method (one-to-one versus canonical correlation), highlighting that dental adaptations to ecological niches are best described by a complex combination of features. The finding of stronger correlations with habitat-related than trophic traits might *prima facie* contradict the role of teeth in food gathering and processing. However, many sharks are opportunistic feeders, completeness of trophic information varies considerably between species, and prey composition may change seasonally and between age cohorts (e.g.^49,50^). Habitats are usually correlated with feeding habits and may serve as a coarser, yet more inclusive, proxy for a species’ primary food sources. Intriguingly, we found striking similarities in heterodonty correlation patterns between specific habitat and food combinations, suggesting discrete clusters representing major strategies: (1) shallow-water habitats and crustacean/small invertebrate diets, (2) open-sea habitats, large preys, with a sub-cluster of deep-sea cephalopod feeders. It is tempting to interpret these clusters in terms of different feeding mechanics^1^. Many shark species specialized on hunting larger vertebrates, which tend to populate in-shore or ocean surface habitats, use their jaws in a saw-like manner^14–16,51^, mechanically benefiting from dental serrations (i.e.fine cusps), absence of larger cusps, and low-to-intermediate heterodonty. Biomechanical studies showed that serrated teeth reduce tear and shear stresses involved in dismembering large prey, while impeding puncturing, reducing suitability for smaller or hard-shelled prey^14–16^. Interestingly, several squalean taxa exhibit dignathic heterodonty with interlocking asymmetric lower teeth and simpler arrow-shaped upper teeth, which cooperate in a holding-sawing mechanism^1^. Consequently, increases in dignathic heterodonty may have enabled the emergence of mechanistically complex feeding strategies especially across Squalomorphii^41,48^. Many benthic feeders, including many deep-sea and reef dwellers, employ complex collecting-crushing or ambushing-grasping strategies aiming at smaller, often hard-shelled, prey, in line with disparate tooth shapes and assemblies along the jaw. Complementarily, feeding strategies often involve jaw-cartilage-level adaptations, indirectly affecting tooth numbers and heterodonty gradients^41,52^. Increased monognathic heterodonty often reflects diversity of specialized functions during different feeding stages. For instance, *Heterodontus* displays different tooth types, reflecting specialization in collecting and crushing hard-shelled prey. Nevertheless, this atypical genus does not commonly stand out in our analyses, suggesting persistent shark-wide trends in heterodonty. Conversely, many smaller reef-dwelling species catch small free-swimming animals using multi-cuspid dentitions^41,42^. Taken together, habitat-heterodonty associations reflect how different environments and prey guilds underlie the dynamic evolution of a finite set of functional feeding strategies with specific signatures both at the tooth and dentition-levels.

We have described two different trends that can, most distinctly, be distinguished by Fourier (tooth-level) complexity and total monognathic heterodonty (i.e.dentition-level complexity), complementarily. Thus, we conclude that species tend to increase either tooth-level or dentition-level complexity, but rarely both of them. Thanks to their moderate association with distinct habitats, these two groups help characterize and interpret key morphological aspects at different scales distinguishing two mechanistically different feeding strategies. Feeding strategies of group one, associated with deep-sea and benthic habitats, involve increased collecting and crushing of hard-shelled prey animals leveraging high individual tooth shape specialization and complexity. Such specializations often involve adaptations of the entire feeding apparatus, i.e.modifications to jaw cartilage shape, articulation and musculature, allowing for suction-based food acquisition mechanisms^41,42^, with possible implications on dental shape. Conversely, the second group uses high-cuspid/serrated, and more homodont, dentitions to catch or dismember swimming prey. Such specialized adaptations require concerted fine-tuning on several levels, thereby strongly selecting against intermediate phenotypes. On the single-tooth level, teeth adapted for specialized diets often occupy extreme positions on morphospaces^38^, in line with this assumption. Thus, the discreteness of the two trends, visualized by divergence within the morphospace, suggest morphological discreteness of highly specialized functional mechanisms. Our analysis shows that the evolution of morphological functionality in a trait featuring repetitive structures, here exemplified by dentitions, involves changes in complexity across scales. This is a common observation among “serially homologous” traits^53–55^, whose units share developmental mechanisms that, during evolution, may accumulate divergent features ultimately leading to individualization. Examples include limbs, vertebrae, and different ectodermal appendages across vertebrates. Interestingly, such organs often nest repeated sub-structures at multiple levels, e.g.feather branches within feathers within plumage regions, or digits on limbs^56–59^. Within dentitions, teeth are arranged in regular rows, but feature repeated structures themselves, namely cusps and smaller cusplets, making them another great example of serially organized structures^60–62^. Differences between units often stem from local differences in developmental regulation within the tissue background, or respective higher-level, structures^63–65^. This is in line with morphometric analyses of specific shark clades or species, which display rather gradual patterns of dental variation^28,29,31,37^, while only a few species, most conspicuously within Hexanchidae, show relatively discrete shape transitions. Thus, by considering complexity across several nested levels of dental organization in sharks, this study is an important extension from the common focus on a single level of morphological organization. In particular, our data allow to quantify and compare the specific importance that each level of organization has with respect to function and adaptive evolution. Overall, we see that monognathic heterodonties have a significantly stronger correlation with ecological traits than dignathic heterodonty and cuspidity. A straightforward interpretation of these results is that specialized food processing types involve correlated combinations of fine-tuned tooth shapes along the entire jaw, with function predicting overall arrangement of tooth shapes. These realizations may be relevant for biomechanical studies, which often quantify single-tooth performances in piercing, slicing or grinding^14–16^. In the light of our results, it may be important to complement those studies with whole-jaw testing paradigms^66^, in line with the distinction of feeding types based on whole dentitions^1^.

Intriguingly, we do not find any significant correlation between low-to-moderate genetic distances and heterodonty differences, implying absence of strong constraints that prevent closely related taxa from developing divergent heterodonty patterns. A similar observation appears to apply to jaw shape differences, permitting stark divergences within relatively short evolutionary timespans^40^. Conversely, disparity of tooth-level complexity, with the exception of cuspidities, appears to increase with genetic distance, suggesting significant phylogenetic constraints. This suggests that adaptive change tends to involve changes on the level of heterodonty rather than tooth morphology, possibly involving changes in developmental parameters along the jaw.

Theoretical and experimental studies in mammals have demonstrated that both gradual and discrete heterodont tooth shape change can be achieved by a gradual change of developmental parameters^24,67^, making tinkering with odontogenesis in a global rather than local manner a suitable way of generating adaptive phenotypic change. We hypothesize that fine-tuning of individual teeth without affecting their neighbours might be more difficult than altering jaw-level gradients of morphogens or developmental factors, which will impact downstream odontogenesis locally. Additionally, studies considering evolution on multiple traits have shown that even functionally intertwined traits such as upper and lower jaw in cichlids can exhibit independent evolutionary dynamics^68^. In the context of shark dentition, this means that dignathic heterodonty should be evolvable within short timespans, particularly in absence of occlusional constraints. This would render dentition an accessible model of hierarchical developmental modularity underlying a mosaique fashion of evolutionary change in a set of functionally or ontologically connected traits^56^.

Besides functional requirements and evolutionary contingency, differences in the frequency of variational patterns may reflect developmental biases^69^. Given the weaker correlation of heterodonty differences with genetic distance compared to tooth-level traits, and a stronger association of heterodonties and ecological specializations, it is tempting to speculate that evolutionary change tends to developmentally originate from alterations in higher-level cues rather than from individually tinkering with low-level features. Evidence for this hypothesis comes from different lines of research: Developmental studies have revealed the explicit involvement of signal gradients from the jaw mesenchyme in establishing differences among mammalian teeth^65,70^. Leveraging developmental transcriptomics, another recent study showed how evolution in one tooth will indirectly affect the developmental regulation of teeth in other positions^71^. On the micro-to meso-evolutionary level, morphometric correlations between different mammalian tooth types suggest regulation by shared, yet not identical, developmental factors^72^, although the degree of covariation varies across traits and species^73^.

Developmental and ontological nestedness might be a major biological principle^56,74^. It has been argued that the repetitive nesting of modules within generative networks can be considered a general principle of how to generate complex yet diverse outcomes that transcends the domain of biology^75^. Theoretical research has emphasized that modularity will increase both robustness to undesirable variation and evolvability^76–78^, while lines of evo-devo research have shown that functionally connected modules can and often do evolve independently or at different rates^68,79,80^. In conclusion, teeth may be considered a model system to understand how nature adapts to environmental challenges not only by the emergence and fine-tuning of hyper-diverse phenotypic modules^81^, but also by tinkering with their modular embedding in a morphological context over different levels of organization.

## Methods

### Shark dentition data acquisition and processing

We extracted entire lateral tooth outlines from selected shark species published on the j-elasmo database (j-elasmo:^43^; outlines downloaded 06-2022). This database contains displays of entire erupted toothrows of over 100 species. The selection for this study was based on the criteria of phylogenetic representativity (i.e.sampling from all extant major clades and avoidance of redundancy by sampling among phenotypically similar sister species) and completeness of dentition, aiming at high coverage of all types of dentitions among extant shark species. Dentitions of which too many teeth overlapped visually were not used. However, negligibly overlapping teeth, i.e. teeth whose partial overlap with adjacent teeth was minor and did not obstruct important features such as cusps, were reconstructed by interpolation and comparison to neighboring teeth and included. More substantially visually obstructed or damaged teeth were excluded from the analysis. Occasionally visible minor damages such as small holes to the enameloid were manually corrected. The exception to the completeness criterion was *Pseudotriakis microdon* which features extremely high counts of relatively small teeth. From this species, only a subset of teeth from different jaw positions was used. In addition, we excluded planctivorous sharks due to their highly specialized dentitions. As sharks keep generating teeth continuously, we defined toothrows as the contiguous sequences of fully erupted teeth along the jaw, from meso/anterior to distal/posterior positions. Due to bilateral symmetry, only one jaw hemisphere was used. As the vast majority of shark teeth are blade-shaped and only feature negligible morphological information in bucco-lingual direction, we decided that 2D lateral views suffice for the purposes of our study. Tooth shape extraction was performed using custom-made tools that automatically identify tooth boundaries and manual segmentation where necessary. Tooth size was not taken into consideration within the scope of this study, because it cannot be explicitly included in the set of shape difference descriptors we used. In other words, size differences can be considered a further, independent, dimension of phenotypic features that may differ between teeth. Since the shape of the basal part of the tooth crown tends to show less morphological-functional fine-tuning, than the upper part that is exposed to nutrition, and at times shows a less defined, undulating or porous transition to the jaw mesenchyme, we decided to only consider the upper dental outline. In preliminary analyses for which the entire tooth outlines were considered, we had found that morphometric patterns were in part driven by the basal rather than the upper part of the tooth crowns. The segmentation point between upper and lower part of the outlines was defined by (1) a visual transition of material, otherwise by (2) the most concave lateral point if a visible inflection could be discerned or (3) the most distant pair of outline points in the lower part. Outline point numbers (1000 per tooth) were equalized by interpolation or data reduction in order to ensure comparability across teeth.

### Ecological information

In order to be able to associate tooth phenotypic information with potential ecological function, we collected proxy features that were widely available in data bases and publications. The trophic level was estimated based on published information from stomach contents or pre-calculated trophic scores as referenced in FISHBASE^82^, shark references^83^, and a number of individual sources (see Sharks_eco_refs.xlsx for references^49,84–104^). In the few cases where suitable trophic information was not available, the phylogenetically closest species for which sufficient records were accessible, were supplied instead. In the occasional case of conflicting values, the more detailed, higher-quality, or more clearly documented of the available sources was used preferentially. In addition, we assigned recorded prey items to larger trophic guilds. Piscivory was not assigned as it is highly unspecific with respect to prey size and trophic level, and because nearly all species include fish into their diet. Another specific diet, planctivory, was omitted, as the three plankton-feeding sharks feature very specialized dentitions, which tend to occupy separate parts within morphospaces^37^. Body length, depth and habitat information was compiled using FISHBASE^82^, Shark references^83^, and Sharks of the World^105^. Extreme body length values were neglected as exceptions, or possibly overstated reports, and the documented ranges for mature male and female individuals were used. Unless noted differently, length values used in our analyses were calculated as the average between upper and lower range limits for females and males, respectively, and the average of those. Depth was annotated similarly, we averaged between the upper and lower range limits that were reported, not considering exceptional reports. We also noted whether shark species were associated with specific habitat types, as within the set of references used. For this assignment, we searched for habitat descriptor terms in encyclopedic literature (as given above) that were not explicitly mentioned as exceptional occurrence. This way, we avoided the need to define arbitrary limits between complementary ecological descriptors. Taken together, we used the following categories: TROPH: average trophic level, VERT: non-osteichthyan vertebrate prey, CEPH: cephalopod prey, MOLL: other molluscan prey, CRUST: crustacean prey, VERM: further small usually worm-like invertebrate prey, OMNI: degree of omnivory or number of documented food categories, SHORE: occurrence near shore line, SHELF: occurrence along the continental shelf zone, REEF, reef habitat, OCEAN: open sea, DEEP: deep sea, BENT: benthic habitat, NECT: nectic habitat, DEPTH: general depth of occurrence, D_LOW_: lower depth limit, D_HIGH_: upper depth limit, D_RANGE_: difference between D_LOW_ and D_HIGH_, SIZE: body length, as calculated above, S_MIN_, S_MAX_: reported extremes of adult/fertile individuals, S_HATCH_: size at hatching or birth. Albeit potentially biased in multiple ways, this compilation of data represents what is currently known and available in the published literature.

### Phylogenetic analysis

To build the phylogenetic tree, the following, slowly evolving and commonly used genetic markers were selected: COI, cytB, NADH2, the ribosomal 12S, 16S genes (with full sequences of 12S+tRNA-Val+16S where available) and rag-1. The choice was made based on availability and a previously published phylogeny^106^. See the NCBI accession numbers in the Supplementary table List_NCBI_refs.xlsx. The sequences were concatenated in the specified order and aligned with MUSCLE^107^ (through the program Unipro UGENE^108^) with default parameter options. Alignment regions with gap values higher than 95% were trimmed. The phylogenetic tree was finally built using the program PHYLIP^109,110^, generating a neighbour-joining tree. The neighbour-joining method was used herein because it aligns with standard approaches in comparable studies that involve similar genetic datasets, allowing for direct comparability to previously published phylogenetic analyses, and because it provides fast computation even for large sets of data and is appropriate for clustering the species relationships when the genetic data is incomplete or heterogeneous. Kimura’s two-parameter model (K2P) was used to compute a distance matrix. We used a boostrap of 1000, Seed values of 5 and the Majority Rule (extended) as a consensus-type choice were applied. The final tree was visualized using TreeViewer^111^. For two species, corresponding genetic information was not available: *Oxynotus centrina* and *Heterodontus japonicus*. In order to build the phylogenetic tree, *O*.*centrina* was added, integrating the phylogenetic analysis of Straube et al.2015^112^, i.e.including the same relative branch lengths as provided therein, while *H*.*japonicus* was assumed to have a position very close to *H*.*zebra*. Although both species belong to the same genus, we decided to include two specimens of *Heterodontidae* to provide more than one data point for this clade. Where used, taxonomic categories were assigned according to literature. However, commonly established taxonomic families that resulted in paraphyly were not assigned and polyphyletic units were assigned independently. Intermediate taxonomic levels (super-families to infraorders) were determined based on the phylogenetic tree at hand. Tree branch lengths normalized by maximal and minimal distance between species pairs were added up to quantify genetic distances (dG).

### Tooth comparison

In the absence of a methodological gold standard to quantify phenotypic similarity between tooth pairs, we devised a set of six measures, thus capturing different aspects of shape, cf.FIG.1B. These similarity measures are then deployed to quantify heterodonty by calculating average pair-wise distances between teeth according to the respective heterodonty definitions.

Partial Procrustes Alignment: To minimize the part of shape difference attributed to relative placement, we performed an incremental shape rotation by up to +/-*π*/8, a shift along x and y axes by up to 10%, and a size change by up to +/-25%. These ranges were determined as sufficient in precursory tests with a subset of shape comparisons. The initial size difference correction was done in two ways, by normalization by total outline length, and normalization by total tooth area, and the lower resulting distance was kept. Tooth area was defined by the outline and a straight line connecting its start and end points. Both tooth outlines were centered on their centroids. For the first three shape distance measures, the Procrustes-aligned configuration issuing the smallest distance between the outline pairs was then considered their definitive distance.

#### a) Euclidean mean distance (EMD)

For each pair of morphologies, we calculated the mean distance between every point (x,y) along one outline L1 and the physically closest outline point (X,Y) of the other morphology L2, irrespective of its relative position. This procedure was conducted both-ways.

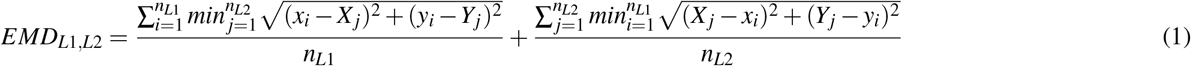

#### b) Homologous Euclidean outline distance (HED)

while the EMD does not make any assumptions about homology, this method compares identical relative positions along two outlines, thus representing pseudo-homology, with distant similarities to semi-landmark methods. i = {1,..,n_L1_} and j = {1,..,n_L2_} are the respective outline points in the two teeth.

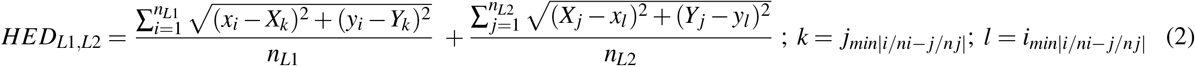

For our specific purposes, n_L1_=n_L2_=n, leading to a simplified formula:

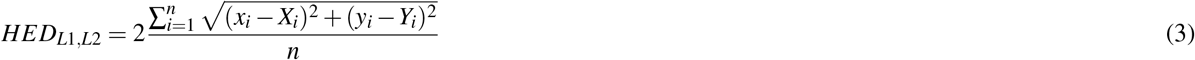

#### c) Superimposed area overlap (SAO)

This method calculates the ratio between the counts of overlapping and non-overlapping parts of the overlaid tooth shapes. The lower boundary delineating the area is defined by a straight line connecting the start and end points of the outlines. To calculate area overlap, both shapes were rasterized into a number of small squares S (i.e.pixels). Before applying Partial Procrustes Alignment, the maximum x and y distances were used to discretize both axes into 100 units, respectively.

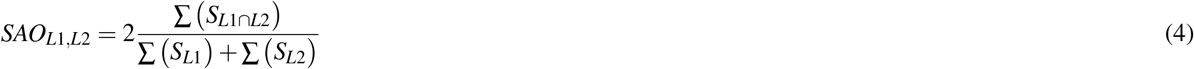

#### d) Discrete Cosine Fourier distance (DFD)

Similar shapes are expected to be defined by sets of similar Fourier coefficients. For this measure, we applied discrete cosine Fourier transformation on the tooth outlines, which incrementally approximates semi-outlines by superimposing cosine lines. Distance is then calculated as the Euclidean distance between all coefficients (for the first 24 harmonics)

Morphological distances between two shapes i and j were quantified by calculating Euclidian distances between the values *z*(*ε*) of the Fourier coefficients *ε, nε* being the number of coefficients at 24 harmonics:

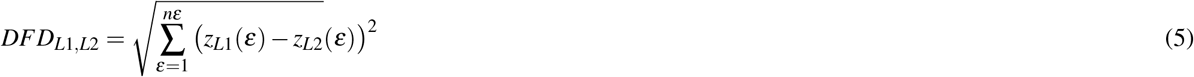

#### e) Outline angle sum distance (OAD)

Outline angles can be calculated between triplets of subsequential outline points. We used the sum function of surface angles for n=100 equidistant outline points (i.e.after point number reduction in order to reduce noise) as a descriptor of shape. We then overlaid the outline angle sum functions *a f* (*i*) of two tooth outlines L1 and L2, starting from the same value, and calculated the area between them. Pairs of similar teeth are expected to show similar functions *a f* (*i*) and low differences in between.

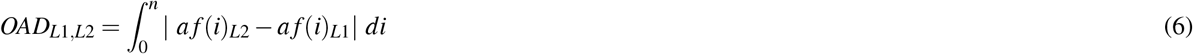

#### f) Angle Function Discrete Cosine Fourier distance (ADD)

In analogy to the DFD, we approximate the specific angle sum function of the OAD by means of discrete cosine Fourier transformation. We then use the Euclidean distance between the resulting Fourier coefficients to describe differences between a given pair of outline functions.

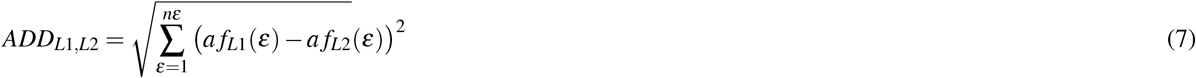

### Heterodonty measures

Heterodonty per dentition was then calculated as the average of all distances, as defined above, between any pair of teeth in consideration. We distinguished between three different heterodonty measures: (a) Sequential monognathic heterodonty (HMS): shape difference between neighboring teeth, (b) total monognathic heterodonty (HMT): shape difference between any pair of teeth within the same jaw, (c) dignathic heterodonty (HDG): shape difference between pairs of teeth at approximately opposite positions on upper and lower jaws. Where the number of teeth differed between the opposing jaws, relative positions were used, eventually causing the same tooth being compared to more than one tooth in the opposing jaw if the latter harbored a larger number of teeth. A schematic of these measures can be seen in FIG.1A. Note that in sharks, no dental occlusion occurs, allowing a higher degree of morphological freedom than in many mammals. In addition, we also recorded the maximum distance between any two teeth within a given jaw (HMX). As a jaw-level descriptor, measures were normalized by division by the respective number of tooth comparisons.

In the following measures, i and j refer to teeth within a respective toothrow or opposing toothrows for the case of dignathic heterodonty.

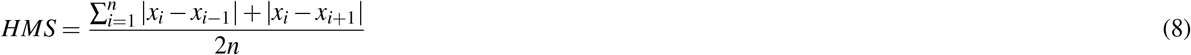

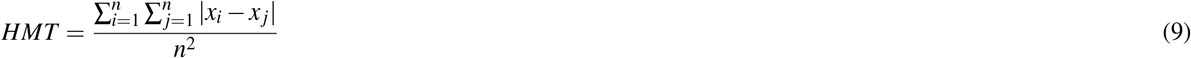

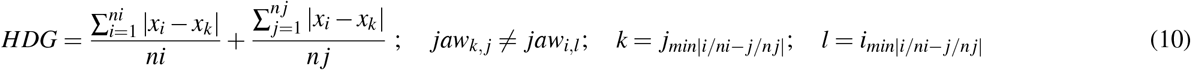

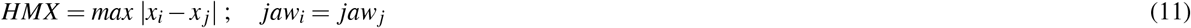

Unless declared otherwise, we calculated the values of heterodonty as the average of all six distance measures devised, in order to minimize potential biases introduced by the choice of a specific method. For inter-species comparison, all values were normalized by the global maxima and minima, respectively.

### Tooth complexity measurements

As for shape distances between pairs of teeth, there is no commonly accepted gold standard method to quantify complexity, even more as it may refer to different features. This is why we devised a range of different methods, as schematically shown in FIG.11. For several analyses, we pooled similar complexity methods, such as outline-based, or angle-based methods. Where not stated otherwise, we calculated total complexity as the sum of all introduced measures. In the displayed formulae, i and j denote outline points of the same tooth, unless explicited otherwise.

a) Coarse-grained cuspidity (CUSP1): the number of larger cusps.
b) Fine-grained cuspidity (CUSP2): the number of minor cusplets. The difference to the previous measure was defined, unavoidably, by an arbitrarily chosen relative size threshold: while the largest cusp was always considered major, cusps were considered minor if their higher col was below 2% of the total length or if they clearly constituted a serration pattern on larger cusps.
c) Outline-to-area ratio (OAR): The total length of the outline was divided by the total area.
d) Outline-to-centroid size ratio (OCR): instead of area, outline length was divided by the centroid size.

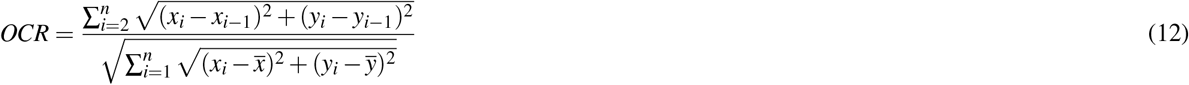

e) Outer/inner circle ratio (OIR): the area of the largest circle inscribed in the outline was divided by the area of the smallest escribed circle encompassing the tooth outline. To prevent a few large ratios from skewing the distribution, we defined a cutoff of 25. This measure captures differences in excentricity.
f) Discrete Cosine Fourier coefficients sum (DFS): due to the definition of Fourier analyses, the size of its coefficients correlates with the excentricity, feature diversity, and difference to a simple round shape. As such, the total sum of Fourier coefficients z for a given outline can be used as a proxy of information required to describe shapes, i.e. shape complexity.

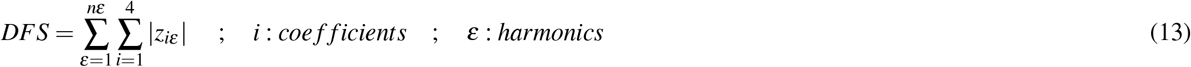

g) Angle sum (ANS): sum of all surface angles (here defined by three points along the outline; angles different from 180° / *π* rad yield higher values.) for NR different resolutions R (defined by the total numbers of equally spaced outline points n_R_). For our angle-based measures, we used six different resolutions with n_R_={8, 16, 32, 64, 125, 250}.

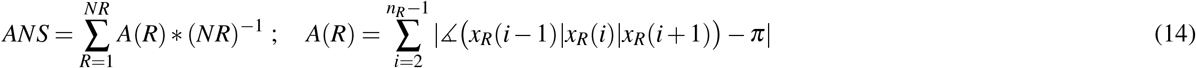

h) Angle sum cadence (ASC): a measure of difference between the angle counts across different resolutions. This reflects the fact that repetitive traits will feature large differences between resolutions, while outlines with differently sized traits will show less difference. In general, the latter case will be considered less complex, as it contains less information.

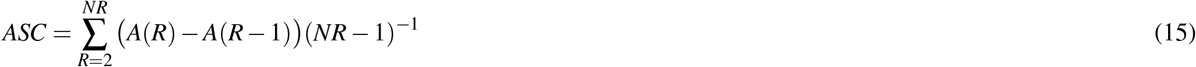
i) Angle disparity (AND): In a similar vein, we measure, for different resolutions, the diversity of angles between pairs of adjacent points on the outlines. Larger diversity is associated with morphological complexity.

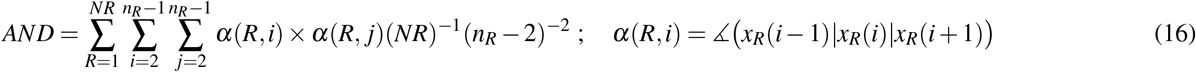

j) Orientation patch count (OPC)^7^: we count the number of contiguous outline streaks that are delimited by a change in absolute direction. Absolute direction is defined by the vectors between subsequent outline points and discretized to absolute partitions of a circle, i.e. a change of direction would correspond to a change of partition and an increase of the count. For this measure, we used different partitions (2,4,8), different rotations of the coordinate system (no rotation, rotation by half a partition and by quarter partitions for the lowest partition number) and the different outline resolutions listed above.

### Phenotypic distance between species

In addition, we calculated total phenotypic distances DP between species pairs (i,j). This is to serve as a test to see how overall similarity would scale with genetic distance, as calculated above. For this measure, teeth of comparable relative jaw positions in two species were compared in a manner analogous to the dignathic heterodonty measure. ni and nj are the total number of teeth per row for the two species, respectively.

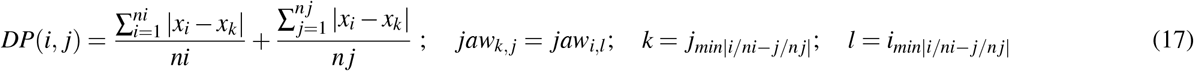

### Statistical analyses

Tooth mean shapes were calculated by averaging the discrete cosine Fourier coefficients within the set of chosen specimens and inversely reconstructing the tooth shape. These operations were performed using the *dfourier* function contained within the R package Momocs^113^. This package was also used to perform shape-based PCA. Canonical correlation analysis based on varying sets of traits was conducted using the R package Cancor. In order to quantify the phylogenetic signal, we took advantage of several of the most frequently used methods: Abouheif’s c-mean, Moran’s P, Pagel’s Lambda, Blomberg’s K. We used available R packages to conduct the analyses: abouheif.moran, moran.idx from the adephylo library and phylosig from the phytools library. We used common R functions (cor, t.test, wilcox.test) as well as the linear regression function via STATS of gnuplot in order to calculate correlation coefficients and p-values.

### Data Visualizations

We used gnuplot (version 5.2) and R (plot and ggplot2 functions) to plot data.

## Supporting information

Sharks_eco_refs.xlsx

List_NCBI_refs.xlsx

## Acknowledgements

The authors are grateful to Fumio Nakagawa for his role in building the j-elasmo database as well as for helpful interactions, and to Arthur Gairin-Calvo, Fidji Berio, Miguel Brun-Usan and Samuel Ginot for critical and constructive comments. This work was supported by a French ANR grant (ANR-21-CE02-0015 PLASTICiTEETH to N.G.) and a Deutsche Forschungsgesellschaft DFG research fellowship (ZI1809/1-1:1, Proj.432922638 to R.Z.).

## Author contributions statement

R.Z. and N.G. designed project; R.Z. collected data and processed images; R.Z. conducted morphometrics and statistical analyses; V.T.-S. performed phylogenetic analysis; R.Z. and V.T.-S. made the figures; R.Z. and N.G. wrote the paper. All authors reviewed the manuscript.

## Additional information

Source of shark dentitions (external: j-elasmo): http://naka.na.coocan.jp/

Ecological data and references: Sharks_eco_refs.xlsx

Genetic sequences used for the phylogeny: List_NCBI_refs.xlsx

Further information will be made available upon request from the corresponding author.

## References

1. Cooper, J. A., Griffin, J. N., Kindlimann, R. & Pimiento, C. Are shark teeth proxies for functional traits? A framework to infer ecology from the fossil record. J. Fish Biol. 103, 798–814, DOI: 10.1111/jfb.15326 (2023).

2. Jernvall, J., Hunter, J. P. & Fortelius, M. Molar tooth diversity, disparity, and ecology in cenozoic ungulate radiations. Science 274, 1489–1492, DOI: 10.1126/science.274.5292.1489 (1996).

3. Fischer, V. et al. Ecological signal in the size and shape of marine amniote teeth. Proc. Royal Soc. B 289, 20221214 (2022).

4. Eronen, J. T. et al. Ecometrics: the traits that bind the past and present together. Integr. Zool. 5, 88–101 (2010).

5. Segall, M. et al. Armed to the teeth: The underestimated diversity in tooth shape in snakes and its relation to feeding behavior and diet. Ecol. Evol. 13, e10011 (2023).

6. Evans, A. R. & Sanson, G. D. The tooth of perfection: functional and spatial constraints on mammalian tooth shape. Biol. J. Linnean Soc. 78, 173–191 (2003).

7. Evans, A. R., Wilson, G. P., Fortelius, M. & Jernvall, J. High-level similarity of dentitions in carnivorans and rodents. Nature 445, 78–81 (2007).

8. Streelman, J. T. & Albertson, R. C. Evolution of novelty in the cichlid dentition. J. Exp. Zool. Part B: Mol. Dev. Evol. 306, 216–226 (2006).

9. Lafuma, F., Corfe, I. J., Clavel, J. & Di-Poi, N. Multiple evolutionary origins and losses of tooth complexity in squamates. Nat. Commun. 12, 1–13, DOI: 1038/s41467-021-26285-w (2015).

10. Davis, A., Unmack, P., Vari, R. & Betancur-R., R. Herbivory promotes dental disparification and macroevolutionary dynamics in grunters (Teleostei: Terapontidae), a freshwater adaptive radiation. The Am. Nat. 187:3, DOI: 10.1086/684747 (2016).

11. Fulwood, E. L. et al. Reconstructing dietary ecology of extinct strepsirrhines (Primates, Mammalia) with new approaches for characterizing and analyzing tooth shape. Paleobiology 47, 612–631, DOI: 10.1017/pab.2021.9 (2021).

12. Evans, A. R. Shape descriptors as ecometrics in dental ecology. Hystrix 24, 133–140, DOI: 10.4404/hystrix-24.1-6363 (2013).

13. Vermillion, W. A., Polly, P. D., Head, J. J., Eronen, J. T. & Lawing, A. M. Ecometrics: A trait-based approach to paleoclimate and paleoenvironmental reconstruction. Vertebrate Paleobiology Paleoanthropology, Methods Paleoecology 373–394, DOI: 10.1007/978-3-319-94265-0_17 (2018).

14. Whitenack, L. B., Simkins Jr, D. C. & Motta, P. J. Biology meets engineering: the structural mechanics of fossil and extant shark teeth. J. morphology 272, 169–179 (2011).

15. Whitenack, L. B. & Motta, P. J. Performance of shark teeth during puncture and draw: implications for the mechanics of cutting. Biol. J. Linnean Soc. 100, 271–286, DOI: 10.1111/j.1095-8312.2010.01421.x (2010).

16. Frazzetta, T. The mechanics of cutting and the form of shark teeth (Chondrichthyes, Elasmobranchii). Zoomorphology 108, 93–107 (1988).

17. Ballell, A. & Ferrón, H. G. Biomechanical insights into the dentition of megatooth sharks (Lamniformes: Otodontidae). Sci. Reports 11, 1232 (2021).

18. Shimada, K. Types of tooth sets in the fossil record of sharks, and comments on reconstructing dentitions of extinct sharks. J. Foss. Res. 38, 141–145 (2005).

19. Maisey, J. G., Turner, S., Naylor, G. J. & Miller, R. F. Dental patterning in the earliest sharks: Implications for tooth evolution. J. Morphol. 275, 586–596, DOI: 10.1002/jmor.20242 (2013).

20. Debiais-Thibaud, M. et al. Tooth and scale morphogenesis in shark: an alternative process to the mammalian enamel knot system. BMC Evol. Biol. 15, 1–17 (2015).

21. Rasch, L. J. et al. An ancient dental gene set governs development and continuous regeneration of teeth in sharks. Dev. Biol. 415, 347–370 (2016).

22. Thiery, A. P., Standing, A. S., Cooper, R. L. & Fraser, G. J. An epithelial signalling centre in sharks supports homology of tooth morphogenesis in vertebrates. Elife 11, e73173 (2022).

23. Zimm, R., Berio, F., Debiais-Thibaud, M. & Goudemand, N. A shark-inspired general model of tooth morphogenesis unveils developmental asymmetries in phenotype transitions. Proc. Natl. Acad. Sci. 120, e2216959120 (2023).

24. Salazar-Ciudad, I. & Jernvall, J. A computational model of teeth and the developmental origins of morphological variation. Nature 464, 583–586 (2010).

25. Harjunmaa, E. et al. Replaying evolutionary transitions from the dental fossil record. Nature 512, 44–48, DOI: 10.1038/nature13613 (2014).

26. Mitsiadis, T. A. & Smith, M. M. How do genes make teeth to order through development? J. Exp. Zool. Part B: Mol. Dev. Evol. 306B, 177–182, DOI: 10.1002/jez.b.21104 (2006).

27. Sharpe, P. T. Homeobox genes in initiation and shape of teeth during development in mammalian embryos. Dev. Funct. Evol. Teeth 3–12, DOI: 10.1017/cbo9780511542626.001 (2000).

28. Shimada, K. Dental homologies in lamniform sharks (Chondrichthyes: Elasmobranchii). J. Morphol. 251, 38–72 (2002).

29. Türtscher, J. et al. Heterodonty and ontogenetic shift dynamics in the dentition of the tiger shark Galeocerdo cuvier (Chondrichthyes, Galeocerdidae). J. Anat. 241, 372–392, DOI: 10.1111/joa.13668 (2022).

30. Tapanila, L., Pruitt, J., Wilga, C. D. & Pradel, A. Saws, scissors, and sharks: Late paleozoic experimentation with symphyseal dentition. The Anat. Rec. 303, 363–376, DOI: 10.1002/ar.24046 (2018).

31. Berio, F., Evin, A., Goudemand, N. & Debiais-Thibaud, M. The intraspecific diversity of tooth morphology in the large-spotted catshark Scyliorhinus stellaris: insights into the ontogenetic cues driving sexual dimorphism. J. Anat. 237, 960–978, DOI: 10.1111/joa.13257 (2020).

32. French, G. et al. The tooth, the whole tooth and nothing but the tooth: tooth shape and ontogenetic shift dynamics in the white shark Carcharodon carcharias. J. Fish Biol. 91, 1032–1047, DOI: 10.1111/jfb.13396 (2017).

33. Cullen, J. A. & Marshall, C. D. Do sharks exhibit heterodonty by tooth position and over ontogeny? A comparison using elliptic fourier analysis. J. Morphol. 280, 687–700, DOI: 10.1002/jmor.20975 (2019).

34. Purdy, R. & Francis, M. Ontogenetic development of teeth in Lamna nasus (Bonnaterre, 1758) (Chondrichthyes: Lamnidae) and its implications for the study of fossil shark teeth. J. Vetrebrate Paleontol. 27(4), 798–810 (2007).

35. Türtscher, J. et al. Evolution, diversity, and disparity of the tiger shark lineage Galeocerdo in deep time. Paleobiology 47, 574–590, DOI: 10.1017/pab.2021.6 (2021).

36. Bazzi, M., Campione, N. E., Ahlberg, P. E., Blom, H. & Kear, B. P. Tooth morphology elucidates shark evolution across the end-cretaceous mass extinction. PLOS Biol. 19, e3001108, DOI: 10.1371/journal.pbio.3001108 (2021).

37. Bazzi, M., Kear, B. P., Blom, H., Ahlberg, P. E. & Campione, N. E. Static dental disparity and morphological turnover in sharks across the end-cretaceous mass extinction. Curr. Biol. 28, 2607–2615.e3, DOI: 10.1016/j.cub.2018.05.093 (2018).

38. Bazzi, M., Campione, N., Kear, B. P., Pimiento, C. & Ahlberg, P. E. Feeding ecology has shaped the evolution of modern sharks. SSRN Electron. J. DOI: 10.2139/ssrn.3770097 (2021).

39. Goodman, K. et al. Ontogenetic changes in the tooth morphology of bull sharks (Carcharhinus leucas). J. Fish Biol. 101, 1033–1046, DOI: 10.1111/jfb.15170 (2022).

40. López-Romero, F. A. et al. Shark mandible evolution reveals patterns of trophic and habitat-mediated diversification. Commun. Biol. 6, 496 (2023).

41. Moss, S. Feeding mechanisms in sharks. Amer.Zool. 17, 355–364 (1977).

42. Wilga, C., Motta, P. & Sanford, C. Evolution and ecology of feeding in elasmobranchs. Integr. Comp. Biol. 47,1, 55–69, DOI: 10.1093/icb/icm029 (2007).

43. Nakagawa, F. J-elasmo. http://http://naka.na.coocan.jp/. Accessed: 2024-03-30.

44. Janvier, P. Early vertebrates (Oxford University Press, 1996).

45. Berio, F. & Debiais-Thibaud, M. Evolutionary developmental genetics of teeth and odontodes in jawed vertebrates: a perspective from the study of elasmobranchs. J. Fish Biol. 98, 906–918 (2021).

46. Salazar-Ciudad, I. & Marín-Riera, M. Adaptive dynamics under development-based genotype–phenotype maps. Nature 497, 361–364, DOI: 10.1038/nature12142 (2013).

47. Adnet, S. Biometric analysis of the teeth of fossil and recent hexanchid sharks and its taxonomic implications. Acta Paleontol. Polonica 51(3), 477–488 (2006).

48. Underwood, C., Johanson, Z. & Smith, M. M. Cutting blade dentitions in squaliform sharks form by modification of inherited alternate tooth ordering patterns. Royal Soc. open science 3, 160385 (2016).

49. Baremore, I., Murie, D. & Carlson, J. Prey selection by the Atlantic angel shark Squatina dumeril in the northeastern Gulf of Mexico. Bull. Mar. Sci. 82(3), 297–313 (2008).

50. Vögler, R., Milessi, A. C. & Quiñones, R. A. Influence of environmental variables on the distribution of Squatina guggenheim (Chondrichthyes, Squatinidae) in the Argentine–Uruguayan common fishing zone. Fish. Res. 91, 212–221, DOI: 10.1016/j.fishres.2007.11.028 (2008).

51. Smith, M. M. et al. Early development of rostrum saw-teeth in a fossil ray tests classical theories of the evolution of vertebrate dentitions. Proc. Royal Soc. B: Biol. Sci. 282, 20151628, DOI: 10.1098/rspb.2015.1628 (2015).

52. Sadier, A. et al. Bat teeth illuminate the diversification of mammalian tooth classes. Nat. communications 14, 4687 (2023).

53. Bateson, W. Materials for the study of variation. Camb. Univ. Press. (1894).

54. Bateson, W. On numerical variation in teeth, with a discussion of the conception of homology (1892).

55. Wagner, G. The biological homology concept. Annu. Rev. Ecol. Syst. 20, 51–69, DOI: 10.1146/annurev.ecolsys.20.1.51 (1989).

56. Zimm, R. et al. Turing’s turtles all the way down: A conserved role of EDAR in the carapacial ridge suggests a deep homology of prepatterns across ectodermal appendages. The Anat. Rec. 306, 1201–1213 (2023).

57. Haupaix, N. et al. The periodic coloration in birds forms through a prepattern of somite origin. Science 361, eaar4777 (2018).

58. Prum, R. & Brush, A. The evolutionary origin and diversification of feathers. The Q. Rev. Biol. 77, 261–295, DOI: 10.1086/341993 (2002).

59. Young, R. L., Bever, G. S., Wang, Z. & Wagner, G. P. Identity of the avian wing digits: Problems resolved and unsolved. Dev. Dyn. 240, 1042–1053, DOI: 10.1002/dvdy.22595 (2011).

60. Stock, D. The genetic basis of modularity in the development and evolution of the vertebrate dentition. Phil.Trans.R.Soc.Lond.B 356, 1633–1653, DOI: 10.1098/rstb.2001.0917 (2001).

61. Luo, Z.-X., Ji, Q. & Yuan, C.-X. Convergent dental adaptations in pseudo-tribosphenic and tribosphenic mammals. Nature 450, 93–97, DOI: 10.1038/nature06221 (2007).

62. Roth, V. On homology. Biol. J. Linnean Soc. 22, 13–29 (1984).

63. Jernvall, J. & Thesleff, I. Reiterative signaling and patterning during mammalian tooth morphogenesis. Mech. Dev. 92, 19–29, DOI: 10.1016/s0925-4773(99)00322-6 (2000).

64. Tucker, A., Matthews, K. & Sharpe, P. Transformation of tooth type induced by inhibition of BMP signaling. Science 282, 1136–1138 (2020).

65. Tucker, A. & Sharpe, P. Molecular genetics of tooth morphogenesis and patterning: The right shape in the right place. J. Dental Res. 78, 826–834, DOI: 10.1177/00220345990780040201 (1999).

66. Corn, K., Farina, S., Brash, J. & Summers, A. Modelling tooth-prey interactions in sharks: the importance of dynamic testing. R.Soc.open sci. 3:160141, 1–6, DOI: 10.1098/rsos.160141 (2016).

67. Harjunmaa, E. et al. On the difficulty of increasing dental complexity. Nature 483, 324–327, DOI: 10.1038/nature10876 (2012).

68. Ronco, F. & Salzburger, W. Tracing evolutionary decoupling of oral and pharyngeal jaws in cichlid fishes. Evol. Lett. 5, 625–635, DOI: 10.1002/evl3.257 (2021).

69. Smith, J. M. et al. Developmental constraints and evolution: a perspective from the Mountain Lake conference on development and evolution. The Q. Rev. Biol. 60, 265–287 (1985).

70. Kavanagh, K., Evans, A. & Jernvall, J. Predicting evolutionary patterns of mammalian teeth from development. Nature 449, 427–433, DOI: 10.1038/nature06153 (2007).

71. Sémon, M. et al. Phenotypic innovation in one tooth induced concerted developmental evolution in another. DOI: 10.1101/2020.04.22.043422 (2020).

72. Gómez-Robles, A., Martinón-Torres, M., de Castro, J. M. B., Prado-Simón, L. & Arsuaga, J. L. A geometric morphometric analysis of hominin upper premolars. Shape variation and morphological integration. J. Hum. Evol. 61, 688–702, DOI: 10.1016/j.jhevol.2011.09.004 (2011).

73. Laffont, R., Renvoisé, E., Navarro, N., Alibert, P. & Montuire, S. Morphological modularity and assessment of developmental processes within the vole dental row (Microtus arvalis, Arvicolinae, Rodentia). Evol. Dev. 11, 302–311, DOI: 10.1111/j.1525-142x.2009.00332.x (2009).

74. Cantor, M. et al. Nestedness across biological scales. Plos One 12(2), 1–22, DOI: 10.1371/journal.pone.0171691 (2017).

75. Solé, R. & Valverde, S. Evolving complexity: how tinkering shapes cells, software and ecological networks. Philos. Transactions Royal Soc. B: Biol. Sci. 375, 20190325, DOI: 10.1098/rstb.2019.0325 (2020).

76. Klingenberg, C. P. Integration, modules and development: molecules to morphology to evolution. Phenotypic integration: studying ecology evolution complex phenotypes 213–230 (2004).

77. Espinosa-Soto, C. & Wagner, A. Specialization can drive the evolution of modularity. Plos Comput. Biol. 6(3):e1000719, 1–10, DOI: 10.1371/journal.pcbi.1000719 (2010).

78. Wagner, A. Robustness and evolvability: a paradox resolved. Proc. Royal Soc. B: Biol. Sci. 275, 91–100, DOI: 10.1098/rspb.2007.1137 (2007).

79. Watanabe, A. et al. Ecomorphological diversification in squamates from conserved pattern of cranial integration. PNAS DOI: 10.1073/pnas.1820967116 (2019).

80. Felice, R. N., Randau, M. & Goswami, A. A fly in a tube: Macroevolutionary expectations for integrated phenotypes. Evolution 72, 2580–2594, DOI: 10.1111/evo.13608 (2018).

81. Jernvall, J. & Salazar-Ciudad, I. The economy of tinkering mammalian teeth. In Tinkering: The Microevolution of Development: Novartis Foundation Symposium 284, vol. 284, 207–224 (Wiley Online Library, 2006).

82. Froese, R., Pauly, D. et al. Fishbase (2010).

83. Pollerspöck, J.. S. Shark references. https://www.shark-references.com//. Accessed: 2024-07-20.

84. Cortés, E. Standardized diet compositions and trophic levels of sharks. ICES J. Mar. Sci. 56, 707–717, DOI:10.1006/jmsc.1999.0489 (1999).

85. Bizzarro, J. J., Carlisle, A. B., Smith, W. D. & Cortés, E. Diet composition and trophic ecology of northeast Pacific Ocean sharks. Adv. Mar. Biol. Pac. Shark Biol. Res. Conserv. Part A 111–148, DOI: 10.1016/bs.amb.2017.06.001 (2017).

86. Ba, B., Diop, M., Diatta, Y., Justine, D. & Ba, C. Diet of the milk shark, Rhizoprionodon acutus (Chondrichthyes: Carcharhinidae), from the Senegalese coast. J. Appl. Ichthyol. 2013, 1–7, DOI: 10.1111/jai.12156 (2013).

87. Li, Y., Gong, Y., Chen, X., Dai, X. & Zhu, J. Trophic ecology of sharks in the mid-east Pacific ocean inferred from stable isotopes. J. Ocean. Univ. China 13, 278–282, DOI: 10.1007/s11802-014-2071-1 (2013).

88. Stevens, J. D. & Cuthbert, G. J. Observations on the identification and biology of Hemigaleus (Selachii: Carcharhinidae) from Australian waters. Copeia 1983, 487, DOI: 10.2307/1444394 (1983).

89. Yano, K. & Musick, J. Comparison of morphometrics of Atlantic and Pacific specimens of the false catshark, Pseudotriakis microdon, with notes on stomach contents. Copeia 1992(3), 877–886 (1992).

90. Osgood, G. J. & Baum, J. K. Reef sharks: recent advances in ecological understanding to inform conservation. J. Fish Biol. 87, 1489–1523, DOI: 10.1111/jfb.12839 (2015).

91. Barnett, A. J.L.Y.,, Abrantes, K. & Awruch, C. Trophic ecology of an abundant predator and its relationship with fisheries. Mar. Ecol. Prog. Ser. 494, 241–248 (2013).

92. Horie, T. & Tanaka, S. Reproduction and food habits of two species of sawtail catsharks, Galeus eastmani and G. nipponensis, in Suruga bay, Japan. Fish. Sci. 66, 812–825, DOI: 10.1046/j.1444-2906.2000.00133.x (2000).

93. Park, J. M., Baeck, G. W. & Raoult, V. First observation on the diet and feeding strategy of cloudy catshark Scyliorhinus torazame (Tanaka, 1908). Reg. Stud. Mar. Sci. 28, 100596, DOI: 10.1016/j.rsma.2019.100596 (2019).

94. Kamura, S. & Hashimoto, H. The food habits of four species of triakid sharks, Triakis scyllium, Hemitriakis japanica, Mustelus griseus and Mustelus manazo, in the central Seto inland sea, Japan. Fish. Sci. 70, 1019–1035, DOI: 10.1111/j.1444-2906.2004.00902.x (2004).

95. Fergusson, I. K., Graham, K. J. & Compagno, L. J. V. Distribution, abundance and biology of the smalltooth sandtiger shark Odontaspis ferox (Risso, 1810) (Lamniformes: Odontaspididae). Environ. Biol. Fishes 81, 207–228, DOI: 10.1007/s10641-007-9193-x (2007).

96. Kindong, R. et al. All we know about the crocodile shark (Pseudocarcharias kamoharai): Providing information to improve knowledge of this species. J. for Nat. Conserv. 63, 126039, DOI: 10.1016/j.jnc.2021.126039 (2021).

97. Vaudo, J. & Heithaus, M. Dietary niche overlap in a nearshore elasmobranch mesopredator community. Mar. Ecol. Prog. Ser. 425, 247–260, DOI: 10.3354/meps08988 (2011).

98. Huveneers, C., Otway, N. M., Gibbs, S. E. & Harcourt, R. G. Quantitative diet assessment of wobbegong sharks (genus Orectolobus) in New South Wales, Australia. ICES J. Mar. Sci. 64, 1272–1281, DOI: 10.1093/icesjms/fsm111 (2007).

99. Burke, P., Meyer, L., Raoult, V., Huveneers, C. & Williamson, J. Multi-disciplinary approach identifies pelagic nutrient linkage by sawsharks. Rev Fish Biol Fish. DOI: 10.1007/s11160-024-09888-6 (2024).

100. Dunn, M., Stevens, D. J.S.F., & Connell, A. Trophic interactions and distribution of some squaliforme sharks, including new diet descriptions for Deania calcea and Squalus acanthias. Plos One 8:3:e59938, DOI: 10.1111/jai.12156 (2013).

101. Carlisle, A. et al. Integrating multiple chemical tracers to elucidate the diet and habitat of cookiecutter sharks. Sci. Reports 2021:11:11809, DOI: 10.1038/s41598-021-89903-z (2021).

102. Yano, K., Mochizuki, K., Tsukada, O. & Suzuki, K. Further description and notes of natural history of the viper dogfish, Trigonognathus kabeyai from the Kumano-nada sea and the Ogasawara islands, Japan (Chondrichthyes: Etmopteridae). Ichthyol. Res. 50, 251–258, DOI: 10.1007/s10228-003-0165-7 (2003).

103. Baremore, I., Murie, D. & Carlson, J. Seasonal and size-related differences in diet of the Atlantic angel shark Squatina dumeril in the northeastern Gulf of Mexico. Aquatic Biol. 8, 125–136, DOI: 10.3354/ab00214 (2010).

104. Campagno, L. J. Alternative life-history styles of cartilaginous fishes in time and space. Environ. Biol. Fishes 28, 33–75 (1990).

105. Campagno, L., Dando, M. & Fowler, S. Sharks of the World. DOI: 10.2307/j.ctv1574pqp (2005).

106. Vélez-Zuazo, X. & Agnarsson, I. Shark tales: a molecular species-level phylogeny of sharks (Selachimorpha, Chondrichthyes). Mol. phylogenetics evolution 58, 207–217 (2011).

107. Edgar, R. C. Muscle: multiple sequence alignment with high accuracy and high throughput. Nucleic acids research 32, 1792–1797 (2004).

108. Okonechnikov, K., Golosova, O., Fursov, M. & Team, U. Unipro ugene: a unified bioinformatics toolkit. Bioinformatics 28, 1166–1167 (2012).

109. Baum, B. R. Phylip: phylogeny inference package. Version 3.2 (1989).

110. Revell, L. J. & Chamberlain, S. A. Rphylip: an r interface for phylip. Methods Ecol. Evol. 5, 976–981 (2014).

111. Bianchini, G. & Sánchez-Baracaldo, P. Treeviewer: Flexible, modular software to visualise and manipulate phylogenetic trees. Ecol. Evol. 14, e10873 (2024).

112. Straube, N., Li, C., Claes, J. M., Corrigan, S. & Naylor, G. J. P. Molecular phylogeny of Squaliformes and first occurrence of bioluminescence in sharks. BMC Evol. Biol. 15, DOI: 10.1186/s12862-015-0446-6 (2015).

113. Bonhomme, V., Picq, S., Gaucherel, C. & Claude, J. Momocs: Outline analysis using R. J. Stat. Softw. 13, 1–24 (2014).

